# Landscape and mutational dynamics of G-quadruplexes in the complete human genome and in haplotypes of diverse ancestry

**DOI:** 10.1101/2025.06.17.660256

**Authors:** Nikol Chantzi, Shiau Wei Liew, Aurell Wijaya, Candace Chan, Ioannis Mouratidis, Emilyane de Oliveira Santana Amaral, Yasin Uzun, Martin Hemberg, Karen M. Vasquez, Chun Kit Kwok, Ilias Georgakopoulos-Soares

## Abstract

G-quadruplexes (G4s) are alternative DNA structures with diverse biological roles, but their examination in highly repetitive parts of the human genome has been hindered by the lack of reliable sequencing technologies. Recent long-read based genome assemblies have enabled their characterization in previously inaccessible parts of the human genome. Here, we examine the topography and genomic instability of potential G4-forming sequences in the gap-less, reference human genome assembly and in 88 haplotypes of diverse ancestry. We report that G4s are highly enriched in specific repetitive regions, including in certain centromeric and pericentromeric repeat types, and in ribosomal DNA arrays, and experimentally validate the most prevalent G4s detected. G4s tend to have lower methylation than expected throughout the human genome and are genomically unstable, showing an excess of all mutation types, including substitutions, insertions and deletions and most prominently structural variants. Finally, we show that G4s are consistently enriched at PRDM9 binding sites, a protein involved in meiotic recombination. Together, our findings establish G4s as dynamic and functionally significant elements of the human genome and highlight new avenues for investigating their contributions to human disease and evolution.

## Introduction

DNA G-quadruplexes (G4s) are alternative DNA structures commonly found in GC-rich areas of a genome. G4s are formed via Hoogsteen hydrogen bonding, which links four guanine bases into a square planar structure known as a G-quartet, which are then stacked on top of each other to form G4 DNA structures (1). The formation of G4s in telomeric regions was among the first indications that these structures represent a naturally occurring DNA conformation (2). Since then, experimental methods have been developed to systematically and in high-throughput detect G4 formation potential (3). The consensus G4 motif, G≥3N1–7G≥3N1–7G≥3N1–7G≥3, has been used to identify sequences predisposed to G4 structure formation and has been implemented in numerous studies (4–6). However, it has become evident that certain G4 structures do not adhere to this consensus G4 motif (7–10). As a result, additional computational methods, such as G4Hunter, have been developed to identify a wider variety of potential G4-forming sequences (11–13).

A variety of studies have shown that DNA and RNA G4s have several functional roles. They are enriched in promoter sequences and are involved in the regulation of gene expression (14–24). G4s can modulate splicing (25–27), can control 3D genome structure organization (28–30), dendrite localization (31), and are regulators of cap-dependent and independent translation (32–38), among other functions. Additionally, G4s are linked to increased genomic instability and are associated with human diseases, including cancer and neurodegenerative disorders (39–44). Although both experimental and computational evidence indicate that G4s are highly susceptible to genomic instability, it has also been proposed that the observed signal may, to a large extent, reflect sequencing errors inherent to short-read sequencing technologies (45).

Additionally, studying G4s in highly repetitive and low complexity regions of the human genome, such as in centromeric, pericentromeric, and sub-telomeric regions, as well as in ribosomal DNA (rDNA) arrays, has been challenging (46). Short-read sequencing technologies have contributed to these challenges because they cannot accurately reveal the sequence composition of these parts of the genome. Such regions are particularly interesting since they are fast evolving and can differ significantly in size and organization in humans (47). Additionally, these loci have been linked to multiple human diseases (48), necessitating further in-depth research. Recent international consortia have utilized long-read sequencing technologies to decode low complexity regions of multiple eukaryotic organisms (49, 50). Importantly, the recent completion of the gap-less human reference genome through the Telomere-to-Telomere (T2T) Consortium (51), including the characterization of the centromeres, enables the examination of the distribution and frequency of repeat elements in regions of the genome that were only partially annotated previously. A recent study utilized the T2T assembly of the human genome and of other primate genomes and studied the recent evolution of G4s in the primate lineage (52). Another study examined the G4 landscape in the recently assembled Y chromosome (53). Furthermore, the Human Pangenome Reference Consortium has generated phased, diploid assemblies from genetically diverse individuals (54), which facilitates the study of polymorphisms and diversity in the human population using high-quality diploid genomes.

Here, we examine the topography, variation and mutational dynamics of G4s throughout the human genome using the latest version of the gap-less, reference human genome assembly (CHM13) and 88 haplotypes from the Human Pangenome Reference Consortium (**Figure 1**). We report that G4s are abundant in certain highly repetitive regions of the genome, can be identified in clusters at centromeric, pericentromeric and rDNA repeats, and we experimentally validate the most frequent G4 motifs in these highly repetitive regions. G4s tend to have lower methylation than expected throughout the human genome and we find that they are genomically unstable, having an excess of all mutation categories, but most notably large indels and structural variants. The mutability of G4s is unevenly distributed, with loops exhibiting a relative excess of substitutions and G-runs showing a relative depletion. Finally, we show that G4s are consistently enriched at PRDM9 binding sites, implicating them in meiotic recombination. Together, our findings establish G4s as dynamic and functionally significant elements of the human genome, linking their topography, epigenetic state, mutability, and role in recombination to genome function, evolution and stability.

**Figure 1:**
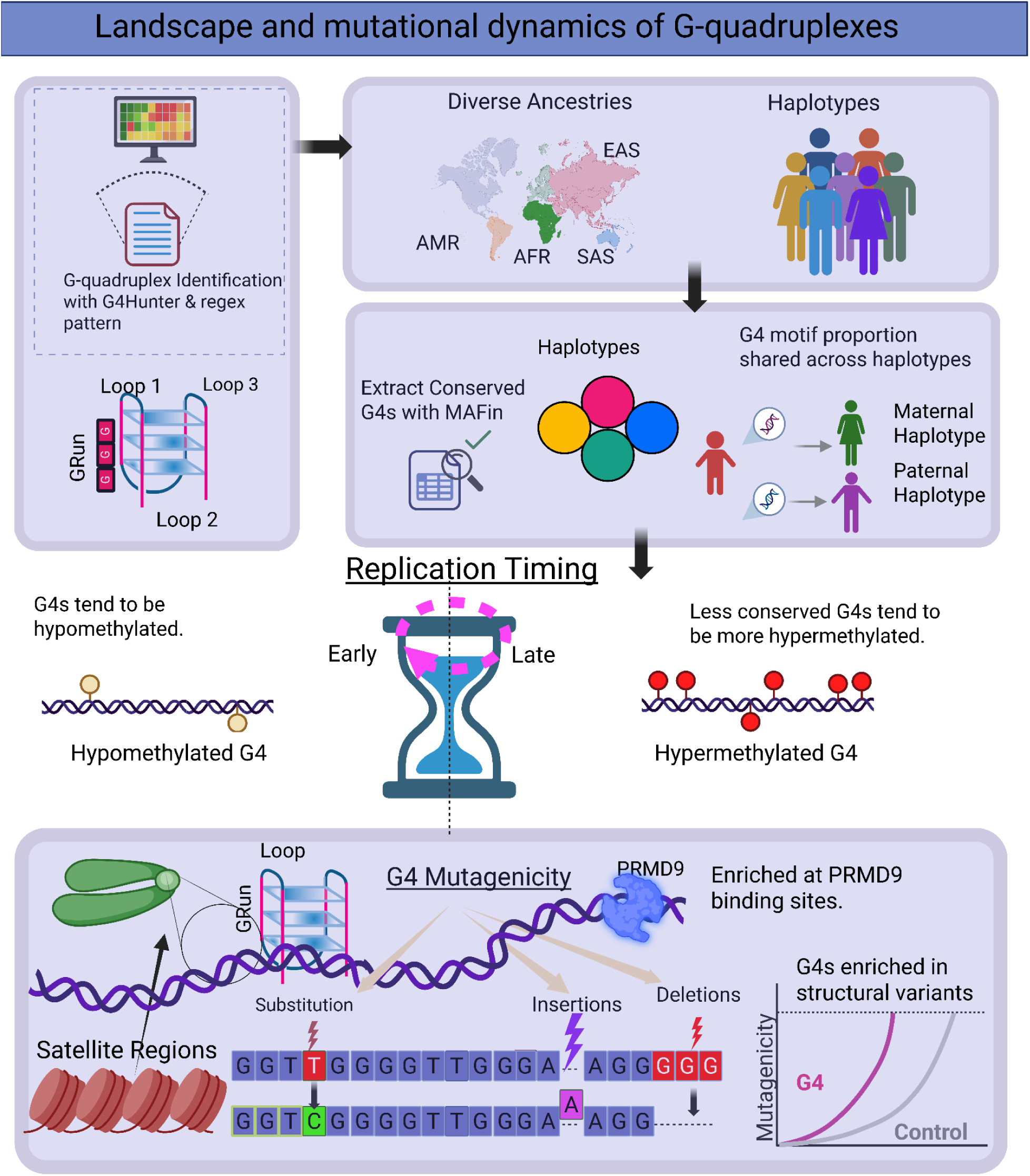
Characterization of G4s across the T2T reference human genome and human haplotypes of diverse ancestry. Schematic illustration summarizing our research process. G4s were identified across the T2T reference human genome and haplotypes of diverse ancestries. Examination of methylation levels, replication timing and germline mutation rate was performed, finding excess mutagenesis and hypomethylation at G4 sites. Examination of highly repetitive elements identified clusters of G4s in rDNA arrays, centromeric transition regions (ct) and Centromeric and Pericentromeric Satellite Annotation (cenSat) sites. PRDM9 binding sites were found to be highly enriched for G4 motifs, in specific PRDM9 genotypes.

## Results

### Characterization of G4s across human haplotypes

Using the recently completed T2T reference human genome (51) and 88 T2T human haplotypes from diverse ancestries provided by the Human Pangenome Reference Consortium (54), we systematically investigated the distribution and population-level conservation of G4s across human chromosomes including in highly repetitive centromeric, pericentromeric and telomeric repeats and ribosomal DNA array loci (47). We identified G4s using two methods, the G4Hunter based algorithm (12) and the regex-based algorithm (55). First, we detected G4s throughout the CHM13 reference human genome assembly and reported a total of 414,904 and 2,081,181 G4s respectively with the two aforementioned methods. These CHM13 reference genome-wide G4 maps were used across the downstream analyses.

We characterized the frequency of G4 sequences and the proportion shared across the 88 human haplotypes. The set of haplotypes included diverse ancestries across four super-populations: African (AFR, n=46), Mixed American (AMR, n=32), South Asian (SAS, n=2), and East Asian (EAS, n=8). Using the G4Hunter algorithm we report an average of 2,048,886 G4s per haplotype and an average of 405,224 for G4Hunter and regex-based methodologies, respectively, with the total number of G4s being relatively stable across the human haplotypes with a standard deviation of 34,233 and 7,608 G4s for G4Hunter and regex-based methodologies, respectively (**Supplementary Figure 1**). We next calculated the total number of unique G4 motifs in each haplotype. Haplotypes from the AFR population tended to contain a higher number of both total and unique G4 sequences compared to other populations. The highest count was observed in a single AFR haplotype, which harbored 20,750 unique G4 sequences identified by the G4Hunter based algorithm and 2,787 by the regex-based algorithm (**Figure 2A**; **Supplementary Figure 1**-**2A**). Our conclusions remained consistent when we combined the G4 motifs from the respective maternal and paternal haplotypes. Genomes of african ancestry exhibited, on average 6,303 and 1,573, unique G4 sequence motifs for G4Hunter and regex-based methodologies, respectively, which were significantly more than any other ancestry group (t-test independent, two-tailed, p-value < 0.001, G4Hunter; p-value < 0.01, regex-based) (**Supplementary Figure 3**).

**Figure 2:**
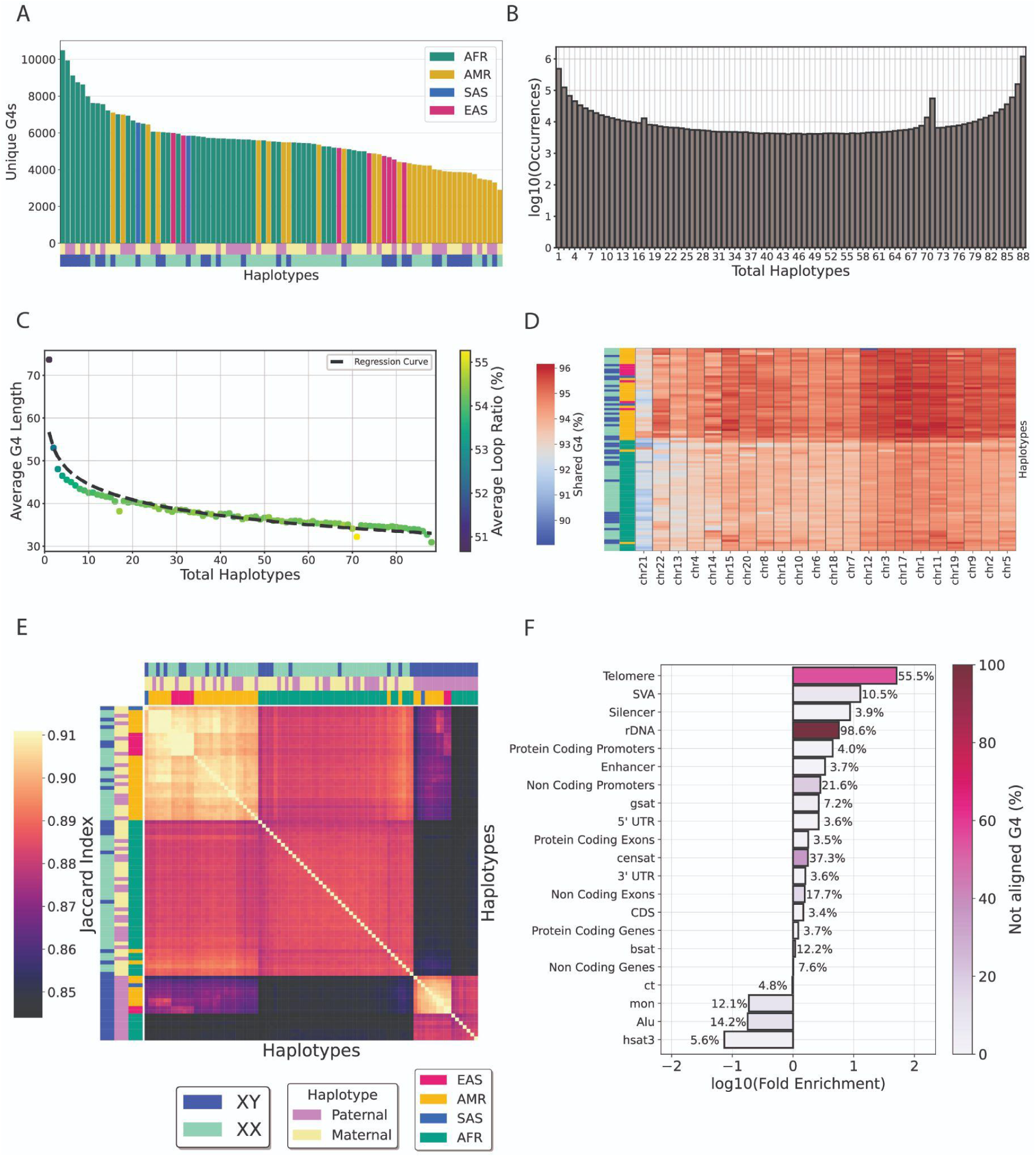
Characterization of G4s across the T2T reference human genome. **A.** Number of unique G4s per human haplotype. **B.** Number of G4s motifs shared across different numbers of haplotypes. **C**. Number of G4 motifs uniquely shared across different numbers of haplotypes plotted against average G4 length. **D.** Shared G4 motifs of each haplotype with the human CHM13v2-T2T reference genome across autosomal chromosomes. **E.** Hierarchical clustering of conserved G4s found in the reference genome CHM13v2 across 88 haplotypes. **F**. Fold-enrichment and ratio of not aligned G4s across haplotypes of G4s across transposable elements, centromeric, pericentromeric and genic subcompartments. Repeats shown include monomeric αSat (mon), classical human satellite 3 (hsat3), γ-satellite (gsat), other centromeric satellites (CenSat), and centromeric transition regions (ct).

We also investigated the extent to which G4 sequences are shared across multiple haplotypes. Using G4s derived from G4Hunter, we observed that 1,199,213 (41.7%) G4 motifs are shared across all 88 haplotypes, while 488,850 (17.0%) are unique to a single haplotype, indicating a distribution favoring shared G4 motifs, followed by haplotype-specific G4s (**Figure 2B**). Similar results were also obtained from the regex-based algorithm (**Supplementary Figure 2B**). When examining the proportion of G4s that are shared across haplotypes, we find that longer G4s are less likely to be found in multiple haplotypes (**Figure 2C**; **Supplementary Figure 2C**), which could provide increased specificity for targeting with G4-based therapeutics (56). We then investigated the proportion of shared G4s across autosomal chromosomes between each haplotype and the CH13v2 T2T human reference genome. Previous studies, showcased that approximately 10% of african pangenome DNA, consisting of 910 individuals, was missing from the GRCh38 human reference assembly (57). We report that haplotypes of AFR ancestry tend to cluster together and share less G4s with the CHM13 T2T reference genome than haplotypes of other ancestries (**Figure 2D**). We calculated the proportion of shared G4 motifs between each pair of haplotypes and found that, as expected, haplotypes from the same population share a higher proportion of G4 motifs and cluster together. However, a distinct cluster emerges due to differences in sex chromosome composition, and an additional subcluster is driven by specific ancestries (**Figure 2E**; **Supplementary Figure 2D**). Thus, we conclude that ancestry plays a role in shaping G4 differences among individuals and that haplotype-specific G4 motifs can be consistently identified.

### G-quadruplexes cluster in the most repetitive parts of the human genome

Leveraging the recent resolution of previously inaccessible repetitive regions of the human genome, we investigated the distribution and haplotype alignment-based conservation of G4s across the 88 haplotypes, including repetitive elements such as pericentromeric and centromeric repeats, as well as ribosomal array regions. We were able to map 92.37% of all G4s identified in the CHM13 reference genome in the multiple alignment file of the human haplotypes. We find that telomeres have the highest G4 density throughout the human genome (**Figure 2F**), which is expected, as the tandem repeats that compose them can fold in G4 structures (2); however G4s in telomeric regions display minimal alignment between haplotypes using the aforementioned conservation thresholds. SVA retrotransposons are found to have the second highest G4 density (12.75-fold enrichment over genome-wide average), which is consistent with the literature (58, 59) and G4s found in SVAs tend to be less conserved among haplotypes. Interestingly, we report a significant enrichment of G4s also at ribosomal DNA arrays, with a 5.68-fold enrichment over background levels, exceeding the G4 frequency at protein coding and non-coding promoters (**Figure 2F**). rDNA and telomeres were the compartments with the least alignable G4s between the haplotypes, followed by Centromeric and Pericentromeric Satellite (CenSat) regions (21.9%). Centromeric transitions (ct) regions while relatively depleted in G4s, displayed high G4 alignability with only 4.8% of G4s not aligning. We conclude that ribosomal DNA arrays are previously overlooked in terms of G4 enrichment, and that G4 conservation between haplotypes varies substantially and is largely driven by the specific subcompartments of the human genome.

We next partitioned each chromosome in equally-sized, mutually exclusive genomic bins of 100kb length, and examined the enrichment and conservation of G4s across haplotypes. To visualize the relationship between G4s and repetitive regions of the genome, we annotated each chromosome with centromeric and pericentromeric satellite repeats, as well as ribosomal DNA array loci. This analysis revealed pronounced differences in G4 distribution both between and within chromosomes (**Figure 3**; **Supplementary Figure 5**). We identified G4 clusters with a 5-fold to over 10-fold enrichment relative to the background G4 density across most chromosomes. In particular, we found that G4s are highly enriched in rDNA regions, and absent from higher order and classic satellite repeats. Of particular interest were the clusters of G4s in the short arm of chromosome 21 (21p) and in chromosomes 13 at rDNA genes (**Figure 3**, **Supplementary Figure 5**). Our results indicate a highly uneven distribution of G4s across chromosomes in the T2T reference human genome, as well as variable chromosome-wide conservation rates across ancestry-diverse haplotypes.

**Figure 3:**
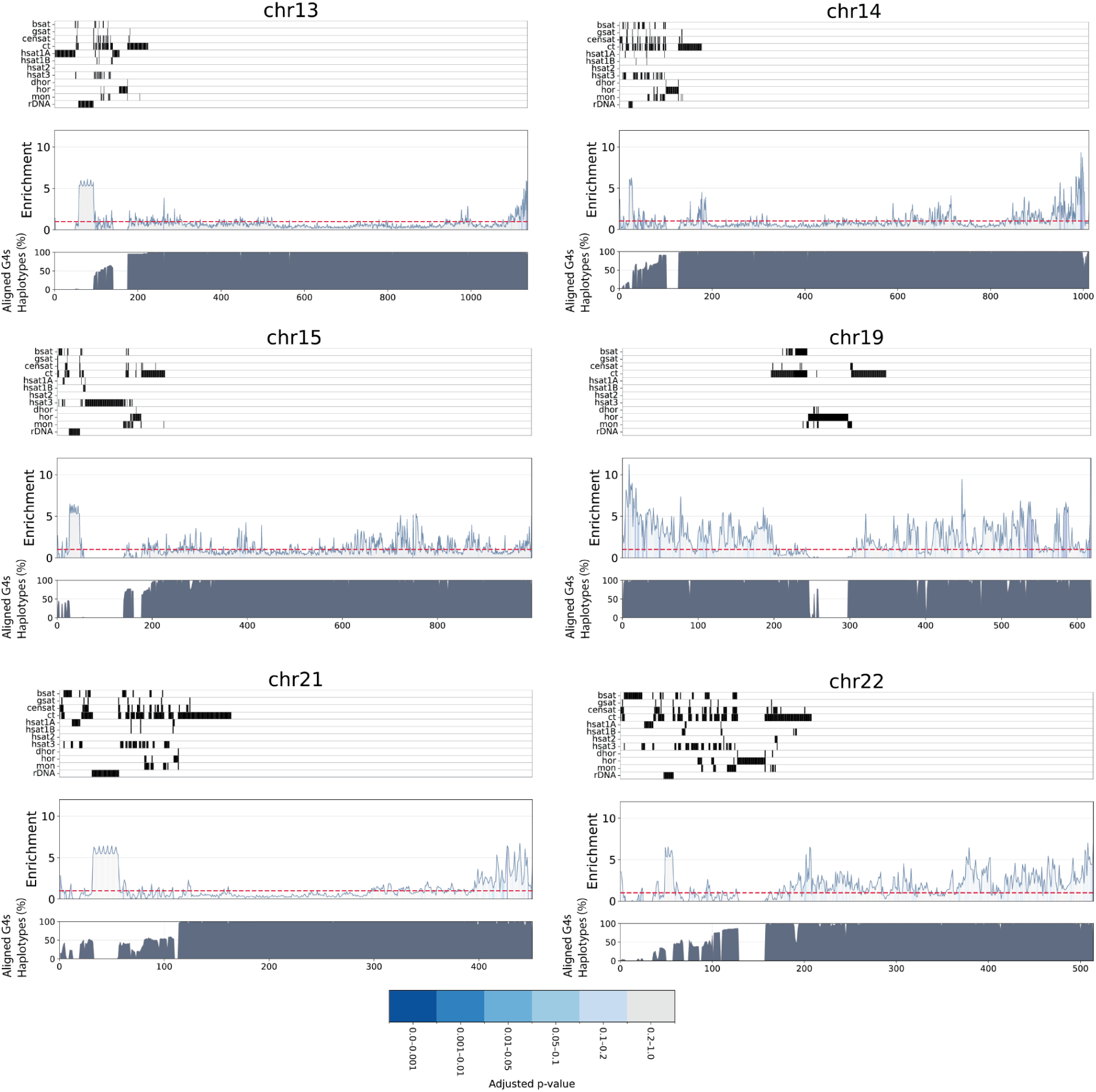
Enrichment of G4s across 13, 14, 15, 19, 21, and 22 human chromosomes, each one equipartitioned into mutually exclusive regions of 100kb length. The area under the curve denotes the empirical p-values (adjusted for multiple comparisons) of the G4 enrichment of each bin, with blue gradient representing a lower p-value, and gray a non-significant enrichment, when adjusting for GC-content. The top panel aligns the position of centromeric and pericentromeric regions. The bottom panel illustrates, for each bin, the average percentage of haplotypes for which aligned G4s were conserved in a given bin. In a given bin, the white stripes represent either the absence of G4s or a low percentage of haplotypes. Repeats include inactive αSat HOR (hor), divergent αSat HOR (dhor), monomeric αSat (mon), classical human satellite 1A (hsat1A), classical human satellite 1B (hsat1B), classical human satellite 2 (hsat2), classical human satellite 3 (hsat3), β-satellite (bsat), γ-satellite (gsat), other centromeric satellites (CenSat), and centromeric transition regions (ct).

### Experimental characterization of G4s in the most repetitive loci of the human genome

Next, we examined the G4 density and sequence composition at highly repetitive regions of the human genome that were only recently resolved in the complete CHM13 reference assembly. Using the G4Hunter algorithm we find that 191,094 G4s, representing 9.18% of G4s in the human genome, fall within centromeric regions. Similarly, we identified 44,010 G4 motifs in centromeres, accounting for 10.61% of all G4s in the human genome, using the regex-based approach. Consistent with our previous findings, rDNA arrays exhibited strong G4 enrichment, showing a 5.7-fold increase over background levels. We detected 35,413 G4 sequences within rDNA, representing 1.7% of all G4s identified by G4Hunter across the complete human T2T genome. Notably, this enrichment surpasses that observed in protein-coding promoter regions, which showed 4.5-fold enrichment. Analogously, for regex-based G4s, rDNA arrays displayed 9.7 fold enrichment with 12,651 G4s located within rDNA arrays, accounting for 3.15% of the total G4s genome-wide (**Supplementary Figure 4**). We found that G4s occupy both genic rRNA and intergenic spacer regions within rDNA arrays. Specifically, genic regex-based G4s are found exclusively in large subunit rRNA, which is consistent with a previous report (60). Among centromeric and pericentromeric repeats, gamma satellite repeats exhibited the highest G4 density, followed by divergent alpha satellite higher-order repeat arrays (**Supplementary Figure 4**). Notably, many of the G4s identified in these repetitive compartments corresponded to highly recurring sequences, suggesting a high level of sequence repetitiveness among G4 motifs in these regions.

We selected a subset of potential G4 DNA-forming sequences, prevalent in centromeric and pericentromeric repeats and in rDNA arrays to experimentally validate their ability to adopt G4 DNA *in vitro*. Interestingly, these were found to occur in certain cases hundreds or even thousands of times within these highly repetitive regions (**Table 1**; **Supplementary Table 1**). For instance, the sequence “GGGTTAGGGTTAGGGTTAGGG” was found 1,957 times in Centromeric Satellite Annotation (CenSat) sites. To determine their G4-forming potential, we performed fluorescent measurements, circular dichroism (CD), and UV-melting spectroscopy experiments. In the fluorescent spectroscopy experiments for both NMM and ISCH-oa1, the fluorescent intensities across all samples were significantly higher in the presence of K^+^ ions compared to Li^+^ ions (**Figure 4A-B**), suggesting G4 structure formation. Additionally, CD spectra revealed higher signal intensities under K^+^ conditions compared to Li^+^ conditions (**Figure 4C**), further supporting the presence of G4 structures. These G4 structures are either in parallel (negative CD peak at 240 nm, positive CD peak at 260 nm) or hybrid (negative CD peak at 240 nm, positive CD peaks at 260 nm and 295 nm) topologies. Furthermore, the UV spectra displayed a hypochromic shift at a wavelength of 295 nm (**Figure 4D**), which is another distinct spectroscopic property to verify the folding of G4 structures in the tested sequences. The maximum negative values observed from the hypochromic shift correspond to their melting temperatures (Tms), which ranged from 50 °C and above (**Figure 4D**), indicating that these G4 DNA structures are thermostable. Overall, the spectroscopic analysis confirmed the formation of thermostable G4 DNA structures under physiological potassium ion concentration within all the selected sequences.

**Figure 4:**
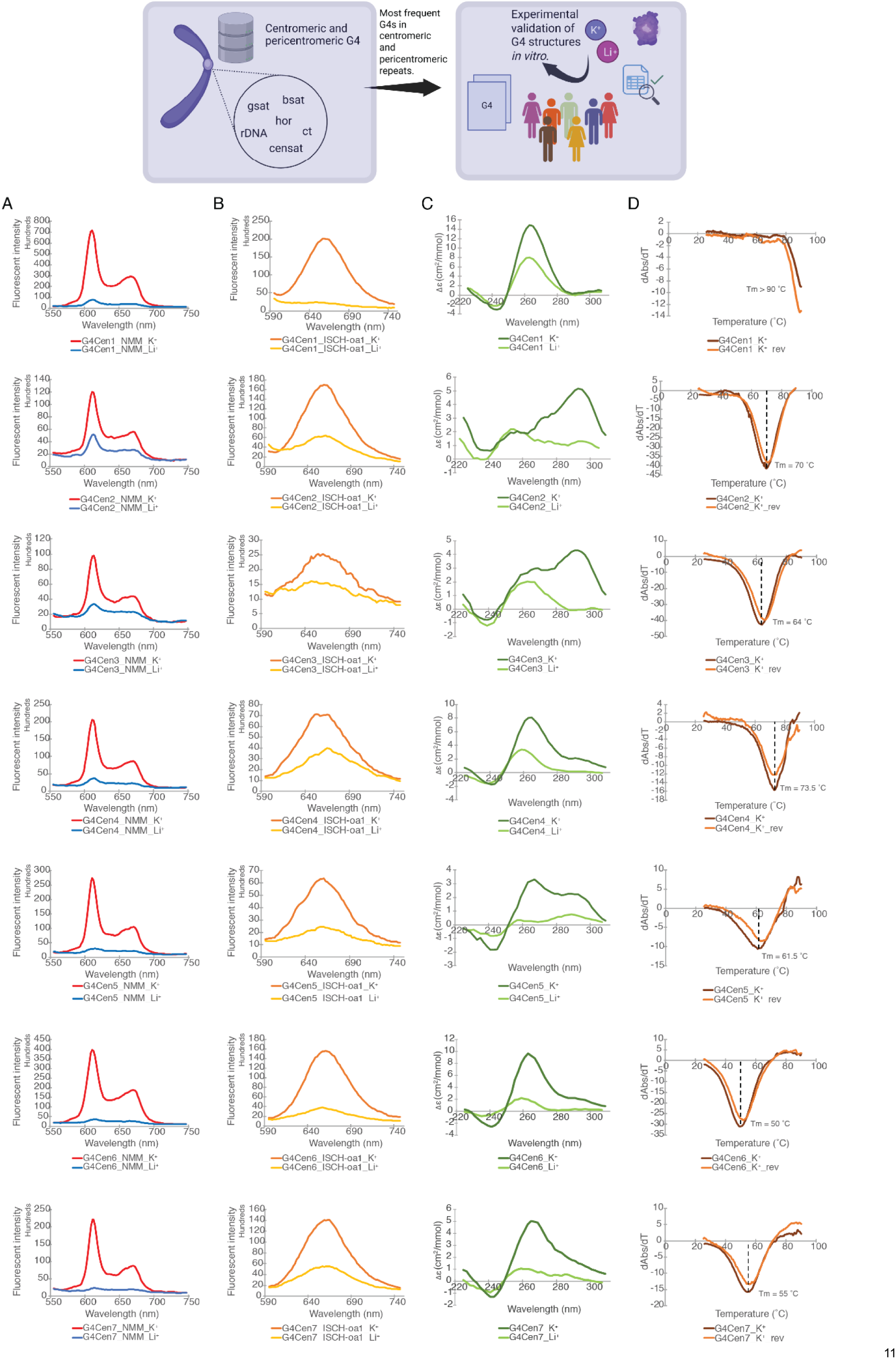
Experimentally validated G-quadruplex candidate sequences. **A)** NMM, and **B)** ISCH-oa1 enhanced fluorescence spectroscopy in the presence of K^+^ and Li^+^. The higher fluorescent signals under K^+^ conditions suggest G4 DNA structure formation in all sequences tested. **C)** CD spectra in the presence of K^+^ and Li^+^. The overall higher signals under K^+^ conditions verify the presence of dG4 in all sequences. The positive and negative peaks at different wavelengths suggest different topologies were formed in different sequences, including parallel (negative peak at 240 nm, positive peak at 260 nm) and hybrid (negative peak at 240 nm, positive peaks at 260 nm and 295 nm) topologies. **D)** UV melting spectra in the presence of K^+^. The hypochromic shift observed at a wavelength of 295 nm is consistent with the presence of DNA G4 structures, with the melting temperature (Tm) determined as the maximum negative value of the hypochromic shift.

**Table 1.**
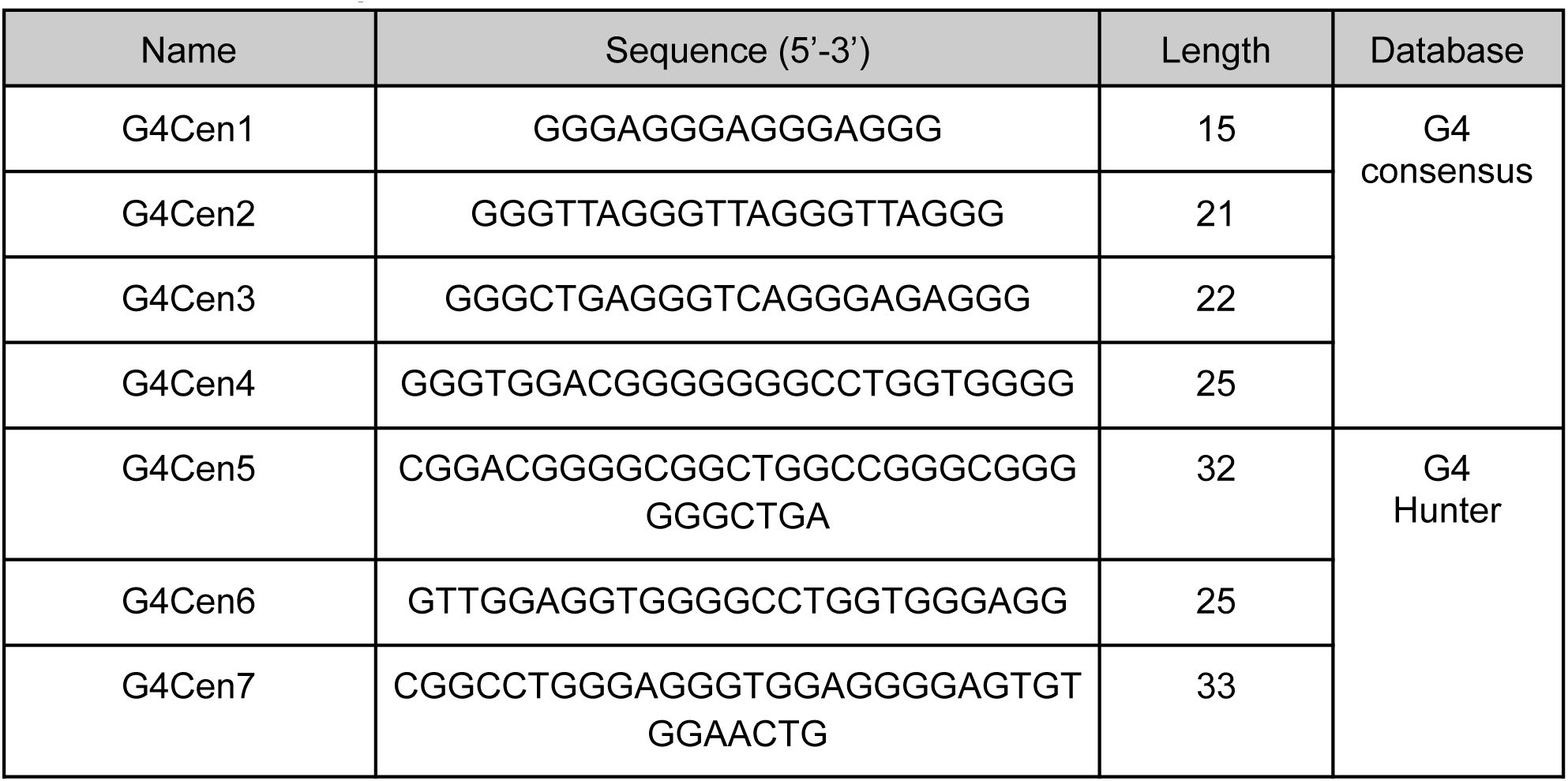
Selected sequences in the centromeres based on G4 consensus and G4Hunter.

### Interplay between the topography and G4 methylation levels

G4s are GC-rich genomic elements and their formation kinetics and stability are influenced by their methylation status (61). Additionally, higher methylation levels have been associated with gene silencing (62). We leveraged the sequenced-based methylation maps generated by ONT nanopore sequencing and HiFi long-read sequencing technologies and examined the effect of the topography of G4s relative to the methylation status across the human genome and within genomic subcompartments. To evaluate if G4s are linked to hyper or hypomethylation we generated G4 controls, adjusting for length, CpG, GpC, and GC content (**Supplementary Figure 6**; Methods), with G4 controls not overlapping any G4 locus.

Overall, we find that G4 motifs exhibit lower average methylation levels than controls, using both the G4Hunter and regex-based extraction methods, in a lymphoblastoid cell line (HG002) and in the CHM13hTERT human cell line (**Figure 5A**; **Supplementary Figure 7**; Mann-Whitney U tests, p-value<0.0001). Next, we utilized Repli-Seq data from an embryonic stem cell line (BG02ES) (63), and examined the methylation status of G4s relative to the replication time decile they were found in. We observe that G4s in early-replicating regions are significantly hypomethylated compared to the matched controls and display greater variability in methylation levels than those in late-replicating regions (**Figure 5B**; Mann-Whitney U, p-value<0.001; Bonferroni adjusted for multiple comparisons). The G4 hypomethylation effect was consistent when we examined the methylation data from the HG002 and CHM13hTERT cell lines and when using G4s derived from the G4Hunter and regex-based algorithms (**Figure 5B**; **Supplementary Figure 8**). It is well-established that CpG methylation is associated with markedly elevated mutation rates (64). Thus, we next examined whether the methylation status of G4s influences their conservation across haplotypes. Notably, in the lymphoblastoid cell line, we identified 884,941 methylated G4s, of which 63,147 had low conservation scores or were not aligned across the studied haplotypes. When comparing methylation levels between conserved and non-conserved G4s, we observed a significant shift toward hypermethylation in the non-conserved group (**Figure 5C**; Mann-Whitney U test, two-tailed, p-value = 0.0). To dissect G4-methylation patterns across the genome, we examined G4s within various genomic subcompartments. Our analysis revealed that in most genic, centromeric, and pericentromeric regions, non-conserved G4s were significantly more likely to be hypermethylated compared to their conserved counterparts (**Figure 5D**; Kolmogorov-Smirnov test, two-tailed, Bonferroni-corrected for multiple comparisons).

**Figure 5:**
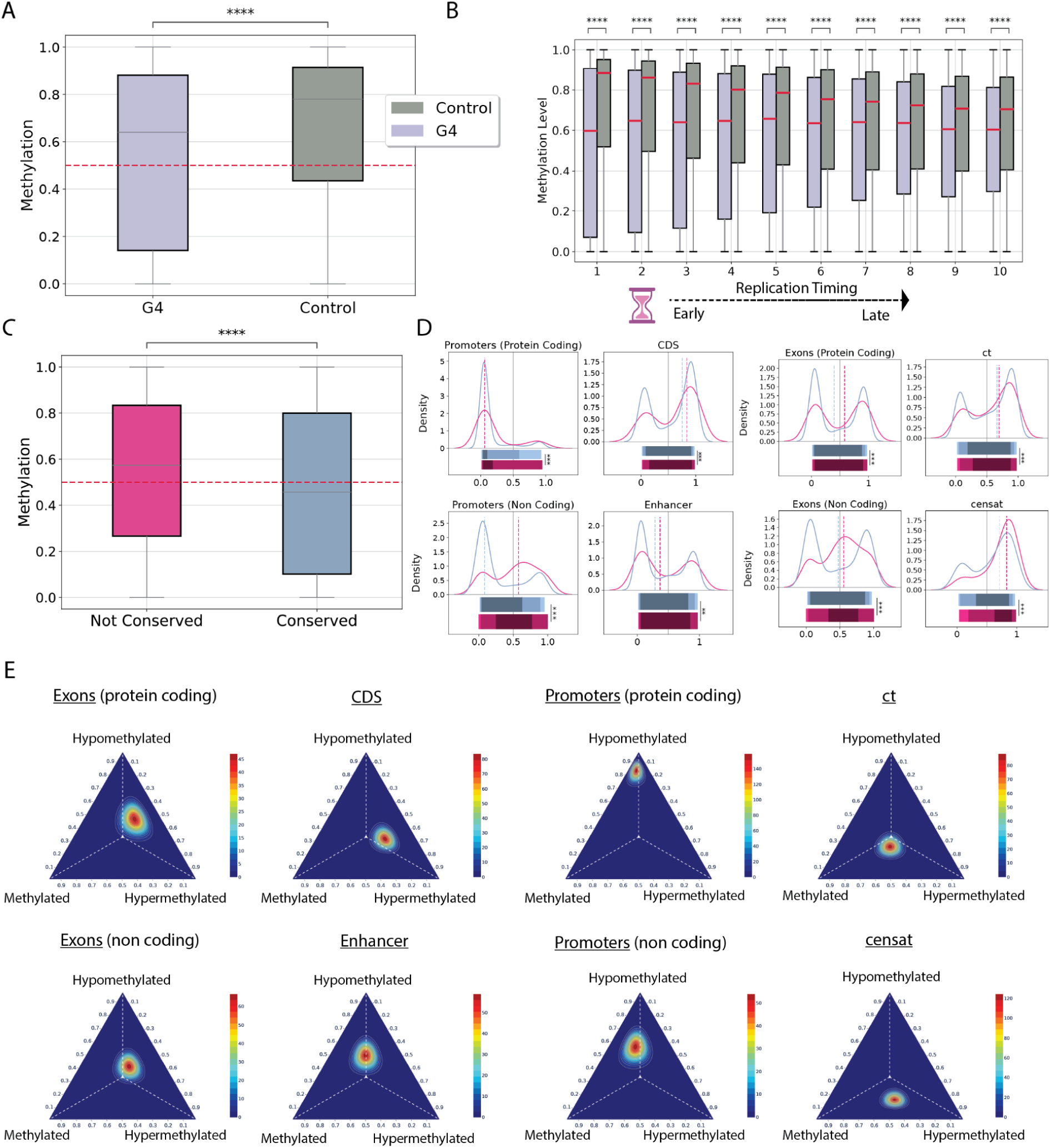
Genome-wide methylation profiles of G4s the complete human genome. **A.** Methylation levels of G4s and their genomic controls using the HG002 cell line. **B.** Methylation levels at G4s and their genomic controls as a function of replication timing domain. **C.** Methylation levels of conserved versus non-conserved G4s. In C-D, statistical significance was assessed using two-tailed Mann–Whitney U tests. **D.** Kernel density estimates comparing methylation profiles of conserved and non-conserved G4s across genomic subcompartments. Statistical significance was evaluated using a two-tailed Kolmogorov-Smirnov test, with Bonferroni correction for multiple comparisons. **E.** Ternary plots illustrating the G4 methylation profiles across various genomic subcompartments. Each profile represents the Dirichlet posterior density. Adjusted p-values are displayed as * for p < 0.05, ** for p < 0.01, and *** for p < 0.001.

As a next step in our analysis we labelled each G4 as hypomethylated, methylated and hypermethylated based on its average methylation status (see Methods). For each genomic subcompartment, we used a maximum likelihood estimator to construct a probability vector representing the distribution of the three G4 methylation profiles. Subsequently, we sampled until convergence from a multinomial distribution and we modeled the resulting counts using a Bayesian Dirichlet distribution. We find that G4s located in protein-coding gene promoters exhibit a strong bias toward hypomethylation, with 82% being hypomethylated. Additionally, G4s found in CDS regions follow a bimodal distribution with higher the probability of hypermethylation. We also analyzed the methylation profile vectors of G4s located in pericentromeric and centromeric regions and found that G4s located in ct regions exhibit a more hypomethylated signal, whereas in CenSat regions the methylation signal is more less hypomethylated (**Figure 5E**). These findings suggest that G4 methylation patterns are shaped by genomic context, with hypomethylation in functional regions potentially contributing to regulatory activity, while hypermethylation in non-conserved loci may reflect reduced selective pressure and epigenetic silencing.

### G-quadruplexes contribute to human genomic instability and polymorphism

Long-read sequencing technologies used for the assembly of CHM13 and for generating the haplotypes of the Human Pangenome Reference Consortium have extremely low sequencing error rates even at highly repetitive parts of the genome (51, 54). We therefore analyzed the mutation rates of G4 sequences across various mutation types, including single-nucleotide substitutions, small insertions and deletions, multi-nucleotide polymorphisms and structural variants (see Methods) from the haplotypes of the Human Pangenome Reference Consortium.

For each mutation site, we centered a 50 base pair window around its midpoint to assess overlap with G4 motifs and quantified G4 fold enrichment relative to the background mutation density. We report an enrichment for germline mutations at G4s across the examined mutation types. More specifically, after adjusting for GC content and correcting for multiple testing, structural insertions (11.27-fold, adjusted p-value < 0.001) and deletions (5.42-fold, adjusted p-value < 0.001) showed the strongest associations, followed by small insertions (3.38-fold, adjusted p-value < 0.0001), small deletions (2.25-fold, adjusted p-value < 0.001), MNPs (2.5-fold, adjusted p-value < 0.01), and substitutions (1.5-fold, adjusted p-value < 0.05) (**Figure 6A**). We examined the relative enrichment of G4s for germline mutations compared to the previously generated G4 controls, which were matched for length, GC content, GpC and CpG content and the results were largely unaltered (**Figure 6B-C**; **Supplementary Figure 9**). We also subdivided the analysis across various genomic regions; we find that G4s consistently exhibited significant enrichment across all mutation loci (**Figure 6D**). These findings indicate that G4s are genomically unstable across mutation categories.

**Figure 6:**
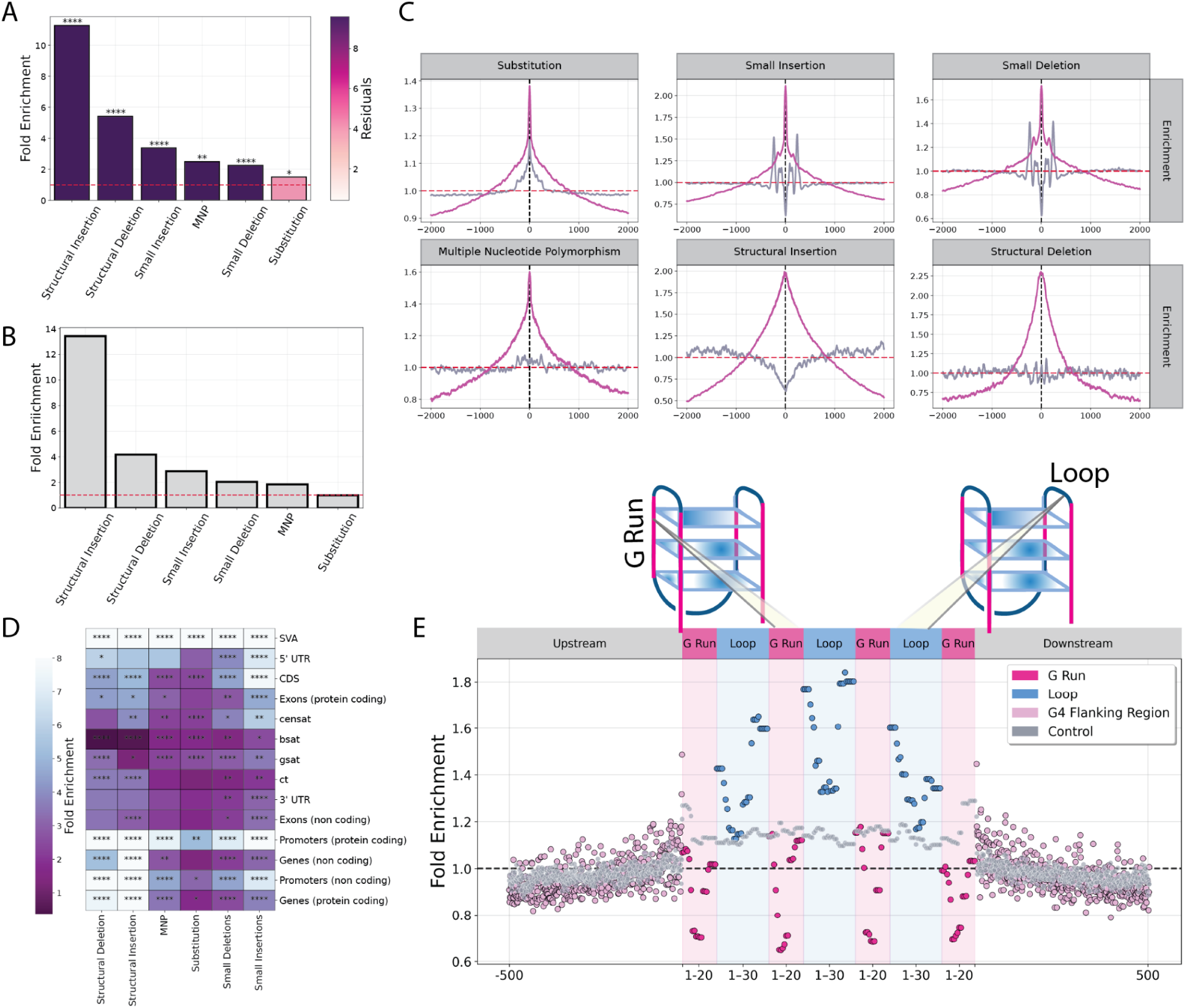
Mutational dynamics of G4s across human genomic compartments. **A.** Fold enrichment of G4 motifs centered at mutation loci expanded into a 100-bp window. Residuals represent the squared deviation between observed and predicted G4 enrichment levels, and p-values indicate statistical significance after correcting for local GC content. **B.** G4-fold enrichment with respect to the control group around the middle of all mutation loci across a 100-bp window. **C.** The distribution of G4s relative to different mutation types including substitutions, small insertions and deletions, multiple nucleotide polymorphisms and structural insertion and deletion breakpoints. A 2-kB window is shown. The two colored lines represent G4s (magenta) and their controls (purple). **D.** Fold-enrichment of mutations at G4s across different mutation categories and different genomic subcompartments. P-values represent the significance of the enrichment after GC-content correction. **E.** Substitution frequency (%) of regex-based G4s in binned G-runs and loops, expanded in a symmetric window 500-bp upstream and downstream. Adjusted p-values are displayed as * for p-value<0.05, ** for p-value<0.01 and *** for p-value<0.001.

We previously reported that the mutability of G4s varies between their subcomponents (39). As the next step in our analysis, we examined mutation events in relation to the G-runs and loops of G4s, focusing specifically on G4s identified using the regex-based algorithm. In G4s with four G-runs, substitutions are concentrated in the center, primarily affecting the central loops and are depleted in the G-runs. This indicates that a substantial proportion of substitutions take place within the G4 loop, especially at the G-run–loop interface (**Figure 6E**). It is established that mutagenicity is strongly determined by the surrounding nucleotide sequence composition (65). To further assess the mutagenicity of G4 loci, we used a trinucleotide composition model, in which we estimated the most frequent mutated trinucleotides across the entire human genome. Subsequently, we estimated the fold enrichment of each mutated trinucleotide within G4 loci relative to the genome-wide average substitution frequency across the window. We found that the most enriched trinucleotides occur at the G-run to loop transition, with GAA being the most enriched (**Supplementary Figure 10**; binomial-test two-tailed, adjusted p-value < 0.001).

Finally, we noticed that 1,378,595 out of the detected 2,065,697 G4 motifs found using G4Hunter, which accounts for the 66,73% of the total G4s in the T2T-CHM13v2, are not overlapping with any mutation type and are conserved across all haplotypes. Analogously, for the regex-based G4s we report that 260,775, which accounts for the 63,4% of the total regex-based G4s detected in the T2T-CHM13v2, are not overlapping with any mutation loci. To investigate each mutation site individually for a specific mutation type within a given genomic subcompartment, we defined the mutation ratio as the proportion of the total number of G4 base pairs overlapping mutation loci to the total G4 sequence length. We report that telomeric regions, SVA transposable elements, centromeric and pericentromeric regions exhibit the highest mutation ratio of G4 base pairs for both G4Hunter and regex-based methodologies (**Supplementary Figure 11**; **Supplementary FIgure 12**). This result was expected as it is consistent with the low conservation reported previously (**Figure 2F**).

### Examination of G4 motifs at PRDM9 binding sites

PRDM9 is a zinc finger protein that mediates DNA double-strand breaks, specifying the positions of recombination hotspots in various mammals, including humans (66, 67). The PRDM9B binding motifs have been previously deduced using chromatin immunoprecipitation with sequencing experiments (ChIP-seq) (68) and PRDM9B binding sites were found to be enriched within rDNA arrays (69). Non-B DNA loci are genomically unstable and are highly enriched for double-strand breaks and we have previously shown that H-DNA motifs, which are are purine:pyrimidine rich mirror-repeat sequences that can form intramolecular triplex DNA, are enriched in PRDM9 hotspots in the human genome (70). Genomic regions have the propensity to adopt multiple non B-DNA conformations under different conditions (71), and H-DNA motifs tend to co-localize with G4 motif sequences. We therefore investigated if PRDM9 binding sites are enriched across G4 sequences.

We examined the genome-wide distribution of G4s relative to PRDM9 ChIP-seq data for different PRDM9 genotypes of different individuals. The experimental data analyzed were derived from normal testis samples from eight individuals. We report that in all eight genotypes analyzed over 50% of PRDM9 binding sites overlap with at least one potential G4 motif (**Figure 7A**). The G4 enrichment at PRDM9 binding sites ranged from 2.94-fold to 1.72-fold in donors AA3 and CL4, respectively (**Figure 7B**). Interestingly, A-like alleles (including A, B, and N alleles) showed stronger enrichments than C-like alleles, and the latter displayed non-significant enrichments when adjusted for GC-content (including C and L4 alleles) (adjusted for multiple comparisons using the Benjamini-Hochberg procedure) (**Figure 7B**). We investigated the relative distribution of G4s within a 2-kb window around PRDM9 loci. We found that G4s exhibited stronger enrichment signals near PRDM9 binding sites compared to the control group, with G4 maximal enrichment peaks proximal to the PRDM9 loci exceeding what would be expected by chance (two-sided Kolmogorov-Smirnov test, *p* < 0.001), with fold changes ranging from 1.48-fold to 1.14-fold in AB1 and CL4, respectively (**Figure 7C**; **Supplementary Figure 13**). Thus, our analysis reveals that G4 motifs are highly enriched at PRDM9 binding sites across multiple human genotypes, with A-like PRDM9 alleles exhibiting stronger G4 overlap than C-like alleles. We conclude that G4s co-localize with PRDM9 binding sites and may play roles in double-strand break formation during meiotic recombination.

**Figure 7:**
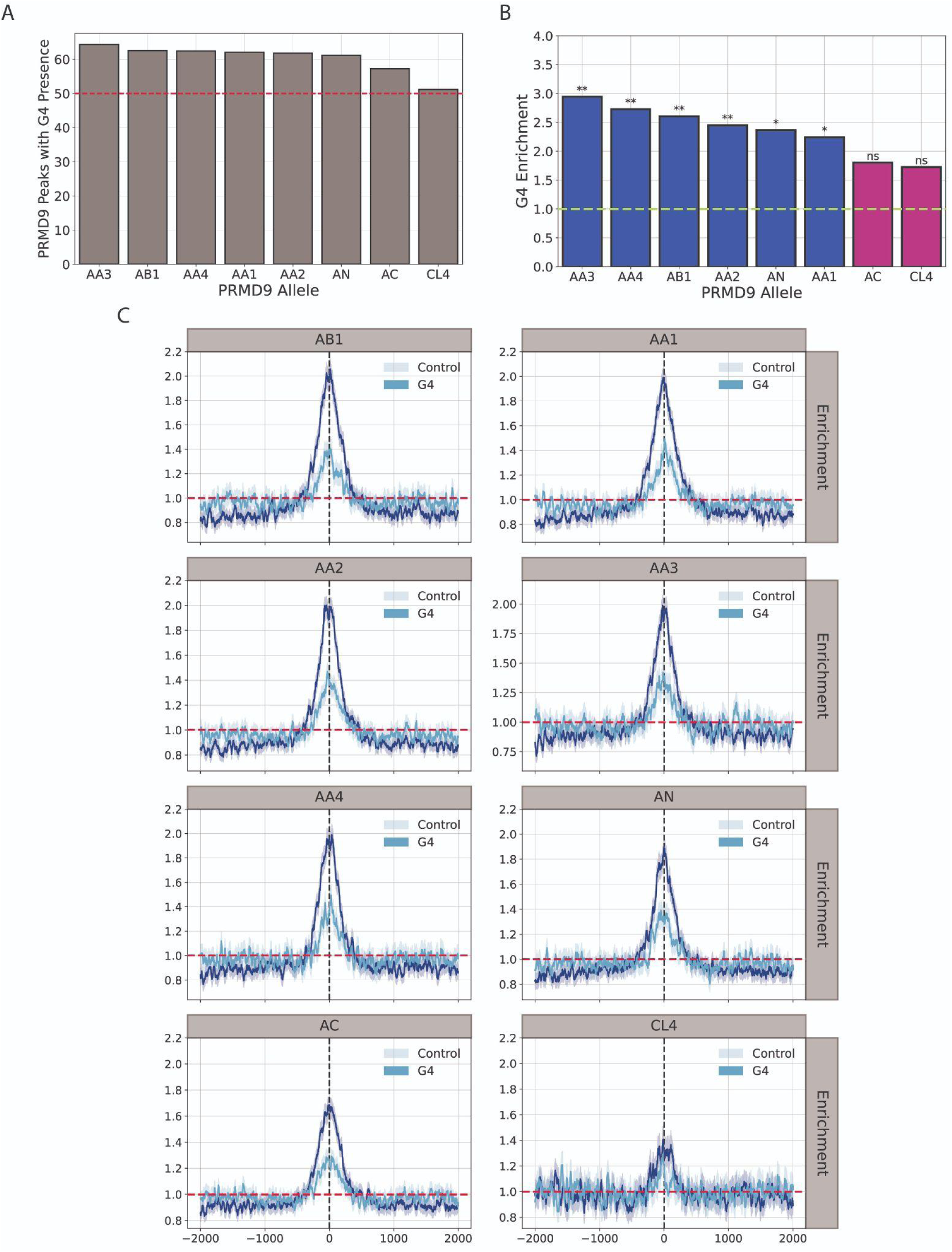
G4 Distribution relative to PRDM9 alleles. **A.** Percentage of PRDM9 regions which contained at least one potential G4 DNA-forming sequence. **B.** GC-corrected enrichment of G4 density relative to the genome-wide G4 density in PRDM9 regions for various alleles. Two-tailed residual tests were performed, and p-values were computed following correction for multiple testing. **C.** Distribution of G4s and the control group, relative to the mid of the PRDM9 region relative to a 2-kB window around the middle. Adjusted p-values are displayed as * for p-value<0.05, ** for p-value<0.01 and *** for p-value<0.001.

## Discussion

Historically, studies have largely focused on G4 DNA in genic and promoter regions, leaving the repetitive landscape of centromeres, pericentromeres, and ribosomal DNA arrays, poorly understood. By leveraging the T2T reference human genome and phased, diploid assemblies from a cohort of genetically diverse individuals representing a human pangenome, we systematically mapped potential G4 DNA-forming sequences across these. Our findings reveal that G4s are unevenly distributed across human chromosomes, are abundant in rDNA arrays and certain pericentromeric repeat elements, such as CenSat, and are notably absent from multiple higher-order satellite repeat types. Interestingly, the G4 enrichment at rDNA arrays exceeds that of protein-coding promoter regions. We also performed multiple biophysical assays and confirmed that G4 motifs in highly repetitive regions of the human genome can indeed adopt G4 structures *in vitro*. Our findings on the enrichment of G4s at rDNA arrays add to previous experimental work that has examined their presence in rRNA (60, 72). However, future work is required to further understand their regulatory roles in ribosomal biogenesis. Additionally, formation of G4s in highly repetitive parts of the human genome could influence genome organization and chromatin structure and *in cellulo* or *in vivo* work is required to further examine if this is the case.

We also find that G4 motifs are hotspots for a wide range of germline mutations, including single nucleotide substitutions, multiple nucleotide polymorphisms, small insertions and deletions, and large structural variants. We find that structural variants are the mutation category most enriched at G4s and are also more likely to be deleterious, as they often disrupt larger genomic regions and thereby increase the likelihood of impacting functional or regulatory elements. Our analysis of G4 subcomponent differences further reveals that for substitutions G-runs and loops within G4s are differentially susceptible to mutation, with loops being particularly vulnerable in accordance with our previous work (39, 73). We conclude that G4s contribute to genetic instability, consistent with previous works (9, 15, 39, 73–75). Such mechanisms are implicated in mutagenesis associated with human diseases including cancer and repeat expansion disorders, underscoring the need to consider G4s in disease models and therapeutic targeting approaches (40, 76). Moreover, the high accuracy of long-read sequencing technologies suggests that the observed G4-associated mutagenesis is unlikely to result from sequencing errors, as previously proposed (45). Au contraire, the conclusions are largely consistent with previous work in germline and somatic mutagenesis indicating that G4s are associated with genomic instability across mutation categories (15, 39).

Our discovery that G4 motifs are highly enriched at PRDM9 binding sites across diverse human genotypes points to a potential functional interplay between non-B DNA structures and the regulation of meiotic recombination (70). Interestingly, A-like PRDM9 alleles showed stronger G4 enrichment than C-like alleles, hinting at allele-specific preferences that could influence hotspot activity. Given that G4s can alter DNA topology and promote double-strand break formation, we propose that G4 structures may facilitate or stabilize PRDM9-mediated recombination events, adding a new dimension to the understanding of recombination hotspot biology.

Our comprehensive analysis reveals that G4s are dynamic, functionally significant elements of the human genome, with highly variable topography, conservation, and mutational profiles across genomic compartments. These findings underscore the importance of G4s in shaping genome stability, recombination, and evolution, including in repetitive regions that were previously inaccessible.

## Methods

### Data retrieval and parsing

The complete human genome (HCHM13) was downloaded from the NCBI database (77, 78). Annotations for centromeres were derived from https://s3-us-west-2.amazonaws.com/human-pangenomics/T2T/browser/CHM13/bbi/censat_v2. 0.bb. Annotations of human functional genomic elements were derived from: https://ftp.ncbi.nlm.nih.gov/genomes/all/GCF/009/914/755/GCF_009914755.1_T2T-CHM13v2.0/ GCF_009914755.1_T2T-CHM13v2.0_genomic.gff.gz. Associated files including genome annotations were downloaded from https://github.com/marbl/t2t-browser. Transposable elements and satellite repeats were annotated using UCSC’s RepeatMasker track (https://hgdownload.soe.ucsc.edu/gbdb/hs1/t2tRepeatMasker/chm13v2.0_rmsk.bb). Methylation data, telomeric regions, and cytogenetic bands annotations were obtained from the Human Pangenome data portal (https://s3-us-west-2.amazonaws.com/human-pangenomics/T2T/CHM13/assemblies/annotation/regulation/chm13v2.0_hg002_CpG_ont_guppy5.0.7_nanopolish0.13.2.bed, https://s3-us-west-2.amazonaws.com/human-pangenomics/T2T/CHM13/assemblies/annotation/chm13v2.0_telomere.bed, https://s3-us-west-2.amazonaws.com/human-pangenomics/T2T/CHM13/assemblies/annotation/chm13v2.0_cytobands_allchrs.bed). We downloaded 88 haplotypes from 44 diploid phased individuals from the Human Pangenome Reference Consortium hosted on the Ensembl genome database: https://projects.ensembl.org/hprc/ from year one data (54). The VCF file of the germline mutations for the human pangenome was obtained from the Human Pangenome Reference Consortium (HPRC) (https://s3-us-west-2.amazonaws.com/human-pangenomics/pangenomes/freeze/freeze1/minigraph-cactus/hprc-v1.1-mc-chm13/hprc-v1.1-mc-chm13.vcfbub.a100k.wave.vcf.gz). For the conservation analysis of G4 loci across the pangenome, a Multiple Alignment Format (MAF) file from the HPRC (https://s3-us-west-2.amazonaws.com/human-pangenomics/pangenomes/freeze/freeze1/minigraph-cactus/hprc-v1.1-mc-chm13/hprc-v1.1-mc-chm13.full.single-copy.maf.gz) was utilized.

### Identification of potential G4 DNA-forming sequences

Regular expressions were employed to generate genome-wide G4 maps using the regular expression of the consensus G4 motif (G ≥ 3N1-7G ≥ 3N1-7G ≥ 3N1-7G ≥ 3) (6). G4s were also detected using G4Hunter with parameters of a window -w =25 and -s 1.5 (12) and processed by a custom Python script to transform the data into a processable tabular format. G4s with overlapping coordinates were merged into a single sequence before performing further analysis.

### Estimation of G4 densities

G4 density was calculated as the number of G4 bps over the number of bps examined. For each chromosome for the T2T reference human genome, we generated mutually-exclusive genomic bins of 100kb equal length. For each genomic bin the G4 fold enrichment was estimated in relation to genome-wide G4 density. We used the residuals of the previously trained second degree polynomial model and the GC proportion of each bin to calculate the predicted G4 enrichment and used a two-tailed test to evaluate the significance of the observed enrichment. Each p-value was adjusted for multiple comparisons within each chromosome using the Benjamini-Hochberg correction. Each aligned G4 from the MAFin (79) output was mapped to a specific bin. Specifically, each start and end coordinate of aligned G4s was assigned to a specific bin, according to the following formula:

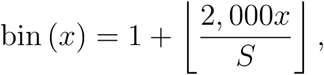

where S denotes the chromosome size. Additionally, we used the bedtools coverage command to estimate for each of the 100kb bins, the G4 enrichment in respect to the genome-wide G4 density. We used MAFin to estimate for each aligned G4, the percentage of haplotypes within each alignment block. For each genomic bin we calculated the average proportion of haplotype presence for aligned G4s as follows:

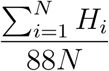

where *H_i_* are the total haplotypes in the MAF block for the *ith* aligned G4, and N denotes total number of aligned G4s within genomic bin G.

We used the scikit-learn Python module to fit the polynomial model using GC content as the predictor variable and G4 fold enrichment as the response. The optimal bin size was selected via grid search with 10-fold cross-validation across the entire binned genome. Finally, for each genomic region, we applied a two-tailed residual test to determine whether the deviation between observed and predicted G4 fold enrichment was statistically significant. Empirical p-values were computed

### Methylation analysis

For each G4 sequence we used “bedtools intersect” to find the total number of methylated locations across each G4 segment (80). To account for multiple methylation values per G4 sequence, we calculated the average methylation level for each G4. The same approach was used to generate an average methylation profile for the control group. We compared the methylation levels of conserved and non-conserved G4s. We classified G4s as non-conserved if they were identified by G4Hunter in CHM13v2 T2T but were not detected in any alignment block or if their alignment score was less than 50%. Subsequently, each G4 segment was classified into a distinct categorical methylation status. Throughout our analysis we classified a genomic segment as hypomethylated if the average methylation level was below 0.2; methylated if the average methylation level was at least 0.2 and lesser than 0.8. Finally, the segment was classified as hypermethylated, if the average methylation level was at least 0.8. This classification is consistent with the literature (81). This classification distributed the G4s to roughly equally partitioned groups.

We utilized Bayesian resampling methods to study the G4 methylation profile in each genomic region. We examined the question that in a fixed genomic compartment if a G4 is methylated what are the odds of being classified as hypomethylated, methylated or hypermethylated. For each genomic subcompartment, we calculated the probability vector representing the fraction of hypomethylated, methylated and hypermethylated G4s, as defined above, found in that region. We used a Dirichlet Distribution with N=3 and uniform prior to model the methylation marks of G4s. We utilized resampling techniques and Bayesian updating to simulate the convergence behavior of the G4 methylation marks in various genomic subcompartments. More specifically, we used the Maximum Likelihood Estimator (MLE) and continuously sampled from a multinomial distribution. At each resampling step we increased the sample size by 1, until the Bayesian estimator, representing the estimated chance for a random methylated G4 having a specific methylation mark, converged to a steady G4 methylation probability vector. For each simulation, we used a Dirichlet joint probability density with conjugate uniform prior to model our initial beliefs about methylation levels, and, at each step, we used the multinomial distribution to sample G4 methylation counts (Hypomethylated, Methylated, Hypermethylated) and the Bayes rule to update the prior distribution. Finally, we used the SciPy library to model the probability density of the resulting posterior Dirichlet distribution and the Plotly library to visualize it as a ternary plot.

### Mutational analysis

The mutation sites were extracted from the VCF file of the Human Pangenome Reference Consortium. We generated an in-house Python script which classifies a given row of the VCF file, by receiving two arguments: the reference allele and the variant(s), and on the basis of their respective lengths, determine the mutation type in the following strictly sequential manner: if the reference allele and the variant have the same length, and the reference allele is of unit length, this is classified as substitution. If the reference allele and the variant have the same length but they are not of unit length, this is classified as multiple nucleotide polymorphism. If the reference length is lesser than the variant length, and the reference length is of length 1 bp, then the mutation is classified as insertion. In particular, the insertion is further classified as small insertion, if the insertion length is lesser than or equal to 50bp; otherwise, it is classified simply as insertion. If the reference length is greater than or equal to the variant length, then the mutation is classified as deletion. Similarly to the insertion case, the deletion is further classified into two distinct categories: small deletion if the deletion length is lesser than or equal to 50bp; otherwise, it’s classified as deletion. The CHM13v2 T2T chromosome Y was excluded from the analysis. Some rows contained multiple variants at the same reference position. To handle this, we parsed each row and, when multiple variants were present, split them into individual entries. Each variant was placed on its own separate row to ensure one variant per row.

### Generation of the Control Group

For each G4 sequence identified by either the G4 Hunter or regex-based method, we generated control sequences with comparable biophysical properties and nucleotide composition. Specifically, for each G4, we randomly selected a position on the same chromosome. From that position, we determined the total length of the original G4 sequence and performed a bidirectional scan—extending both upstream and downstream by the same length—searching for a segment that matched the required biophysical characteristics.

If the segment did not meet our criteria, we iteratively shifted the window by one base pair in both directions while preserving the original sequence length, continuing the search. If no suitable match was found after a predefined number of attempts, the search for that specific control was abandoned, and we proceeded to the next G4. To account for potential biases due to repetitive or AT-rich regions that could hinder the search, we aimed to retrieve two control sequences per G4.

A control sequence was accepted if it matched the G4 in length and fell within a 10% reciprocal tolerance range for GC content as well as CpG and GpC dinucleotide counts. After generating the control sequences, we ensured that none overlapped with any G4 region from either method by using bedtools intersect -v, which filtered out contaminated matches. This careful extraction strategy allowed us to construct a stringent control group with a biophysical profile closely mirroring that of the G4 sequences, particularly with respect to elevated GC content and CpG density.

### GC Content adjustment

We modeled the relationship between GC content and G4 fold enrichment using a second-order polynomial. To do this, we partitioned the CHM13v2 T2T human genome into mutually exclusive bins. For each bin, we computed the fold enrichment of G4s relative to the genome-wide G4 density. This relationship was then modeled using tools from the scikit-learn Python library, with the bin size optimized through grid search and ten-fold cross-validation applied across the entire genome. To assess whether G4 fold enrichment in a given genomic region deviates significantly from expectation, we performed a two-tailed residual test. Fold enrichment for each region was calculated as the ratio of the G4 density in the region to the genome-wide G4 density. We then modeled the relationship between GC content and G4 fold enrichment using a quadratic polynomial, trained on non-overlapping two-megabase windows from the human T2T genome. For each region, we computed the residual, defined as the difference between the observed and expected G4 fold enrichment based on the fitted model. This residual was then used to evaluate statistical significance by determining its percentile within the empirical distribution of residuals from the two-megabase model, using the percentile score function from the SciPy Python library. The resulting empirical p-value was multiplied by two to account for the two-tailed nature of the test. This framework enabled us to identify regions with significantly elevated or depleted G4 enrichment, particularly within functionally distinct genomic compartments such as satellite, centromeric, and pericentromeric regions.

### Mutation density in G-runs and Loops

A G-run is defined as a subsequence of three or more consecutive Gs, while a loop refers to the genomic region between two G-runs. We expanded the regex-based G4 regions using a symmetric window of 500 base pairs upstream and downstream, and used the bedtools intersect -wo command to calculate overlaps with substitutions. Each G4 was split into mutually exclusive G-runs and loops. For this analysis, we retained only canonical G4s containing exactly four G-runs and germline substitutions with allele frequency at least 0.05 or higher. Because most G-runs and loops vary in length, each G-run and loop was normalized into 20 and 30 bins, respectively. We then represented mutation occurrences at each locus using a vector. To homogenize G4s with varying G-run and loop sizes, single base-pair substitutions within G-runs and loops were distributed across multiple bins, whereas substitutions in the flanking regions were assigned to a single bin. Finally, fold enrichment was calculated by dividing the number of mutations at each vector position by the average mutation count across the entire window. The same analysis was performed for the control group, using the corresponding G4 sequence to define the split into G-runs and loops.

### Trinucleotide mutation model

To assess mutagenicity at G4 loci, we constructed a trinucleotide mutation model. We first used Jellyfish (82) to count genome-wide trinucleotide k-mer frequencies in the CHM13v2 T2T human reference genome. Each germline substitution locus was expanded to include its immediate flanking bases. For each of the 32 canonical trinucleotide substitution types, we estimated genome-wide mutagenicity by dividing the number of observed substitutions by the total number of corresponding trinucleotide occurrences in the genome. To evaluate the enrichment of mutagenic trinucleotides within G4 regions, we first used the bedtools merge command from BEDtools (80), to collapse overlapping G4 sequences, then expanded each locus by one base upstream and downstream. We counted the total occurrences of each canonical trinucleotide within these extended G4 regions and used bedtools intersect command to identify which of these overlapped with a substitution locus. The mutagenicity ratio within G4s was calculated similarly to the genome-wide case, and fold enrichment was defined as the ratio of the G4 mutagenicity to the genome-wide mutagenicity for each trinucleotide substitution type.

### Investigation of G4 motif enrichment for different PRDM9 alleles

We analyzed ChipSeq data reported by Alleva et al. (83), for eight individuals with various PRDM9 genotypes, either homozygous (A/A), or heterozygous (A/B, A/N, A/C or C/L4). For each individual, we used liftOver to map the PRDM9 called peaks in hg19 to the T2T-CHM13v2.0 genome. We centered each ChIP-seq peak and created a 2 kb window around the midpoint. After merging overlapping G4 motifs, we used the command bedtools intersect to identify overlaps between the G4s and each window. For each overlap, we counted how often G4 motifs occurred at each position from -2 kb to +2 kb relative to the PRDM9 midpoint. These counts were then normalized by the average across the window to assess G4 enrichment at each position. We calculated 95% confidence intervals using bootstrap sampling (N = 1,000, with replacement) as described above. At each position relative to the PRDM9 midpoint, the lower and upper bounds correspond to the 2.5th and 97.5th percentiles of the enrichment values across bootstrapped samples. The barplot in Fig. 5c shows the maximum G4 enrichment within the 4 kb window centered on the PRDM9 peak. Bars were colored green if the peak enrichment occurred within ±250 bp of the PRDM9 midpoint, and gray if it fell outside this region.

### Identification and conservation analysis of potential G-quadruplex-forming sequences in the pangenome

G4 motifs were first identified in the CHM13 T2T sequence using G4Hunter and the regex-based expression algorithms. Following the identification of the G4 motifs in CHM13 T2T, the corresponding sequences from other pangenome samples were extracted based on their alignment to the CHM13 sequence within the MAF file, ensuring a direct correspondence using MAFin (79). The output file from MAFin was used to determine the alignment score as well as the total number of haplotypes of the alignment block for each aligned G4. We did not take into account the Y chromosome throughout this analysis.

### Verification of the formation of G-quadruplex structures in centromere sequences Oligonucleotides and ligand preparation

The oligonucleotides (oligos) were synthesized by Genewiz Biotechnology Co., Ltd. (China).

They were dissolved in ultrapure nuclease-free distilled water (Thermo, USA) prior to use. The sequences of the oligonucleotides can be found in Table 1. N-methyl mesoporphyrin IX (NMM) was purchased from Frontier Specialty Chemicals (USA), and ISCH-oa1 was received as a gift from Professor Shuo-Bin Chen from Sun Yat-Sen University. NMM and ISCH-oa1 were dissolved in DMSO prior to use. Subsequently, the oligonucleotides and ligands (NMM/ISCH-oa1) were stored at -20 °C before the experiment.

### Fluorescence spectroscopy

Reaction mixtures were prepared by mixing 1 μM oligos with 10 mM LiCac buffer (pH 7.0) and 150 mM KCl/LiCl, reaching a final volume of 100 µl. The mixtures were incubated at 95 °C for 5 minutes to denature the oligos and then left at room temperature for another 10 minutes for renaturation. After that, 1 μM of G4 ligand (NMM or ISCH-oa1) was added to each sample. The fluorescence spectroscopy was performed on the samples in 1 cm path-length quartz cuvettes using a HORIBA FluoroMax-4 fluorescence spectrophotometer. NMM and ISCH-oa1 had their excitation wavelengths set at 394 nm and 570 nm, respectively, and the emission spectra were collected at a 2 nm interval from 550 to 750 nm for NMM and 590 to 750 nm for ISCH-oa1. The entrance and exit slits were set to be 5 and 2 nm for both ligands. All data were blanked and smoothed over 10 nm using Microsoft Excel.

### Circular dichroism (CD) spectroscopy

Reaction mixtures were prepared by mixing 5 μM oligos with 10 mM LiCac buffer (pH 7.0) and 150 mM KCl/LiCl, reaching a final volume of 2 ml. The mixtures were incubated at 95 °C for 5 minutes to denature the oligos and then left at room temperature for another 10 minutes for renaturation. CD spectroscopy was then performed on the samples at a 2 nm interval from 220 to 310 nm using a Jasco CD J-150 spectrometer equipped with 1 cm path-length quartz cuvettes. The scanning was performed using Spectra ManagerTM Suite, and the data were normalized and smoothed over 10 nm using Microsoft Excel.

### UV melting spectroscopy

The samples for UV melting analysis were prepared and denatured as per the CD spectroscopic experiment. UV melting spectroscopy was performed on the samples using a Cary UV-Vis Multicell Peltier spectrophotometer with Teflon tape-sealed 1 cm path-length quartz cuvettes. The UV melting curve was monitored from 20 °C to 95 °C at 0.5 °C intervals at a wavelength of 295 nm. The data were blanked and smoothed over 10 °C using Microsoft Excel.

### Author Contributions

N.C., and I.G.S conceived the study. N.C., wrote the code, generated the visualizations and performed the analyses with help from I.G.S. I.G.S. supervised the project and provided resources. S.W.L. and A.W. performed the G4 experiments with supervision and resources from C.K.K.. N.C, and I.G.S., wrote the manuscript with help from all authors.

### Software availability

The code can be found at https://github.com/Georgakopoulos-Soares-lab/g4_t2t_identification.

### Competing interest statement

The authors declare no competing interests.

### Funding and Acknowledgement

Research reported in this publication was supported by the National Institute of General Medical Sciences of the National Institutes of Health under award number R35GM155468 awarded to I.G.S. S.W.L., A.W., and C.K.K. were supported by National Natural Science Foundation of China (NSFC) Project [32471343 and 3222089] to C.K.K.; Research Grants Council (RGC) of the Hong Kong Special Administrative Region (RFS2425-1S02, CityU 11100123, CityU 11100222, CityU 11100421 to C.K.K.); Croucher Foundation Project (9509003) to C.K.K.; State Key Laboratory of Marine Pollution Seed Collaborative Research Fund (SCRF/0037, SCRF/0040, SCRF0070) to C.K.K.; City University of Hong Kong projects (7030001, 9678302) to C.K.K; and the Hong Kong PhD Fellowship Scheme to S.W.L.; Coordenação de Aperfeiçoamento de Pessoal de Nível Superior - Brasil (CAPES) Finance Code 001 - to E.O.S.A. K.M.V. was supported by the National Cancer Institute of the National Institutes of Health under award number R01CA093729 (to KMV).

## Supplementary Material

**Supplementary Figure 1:**
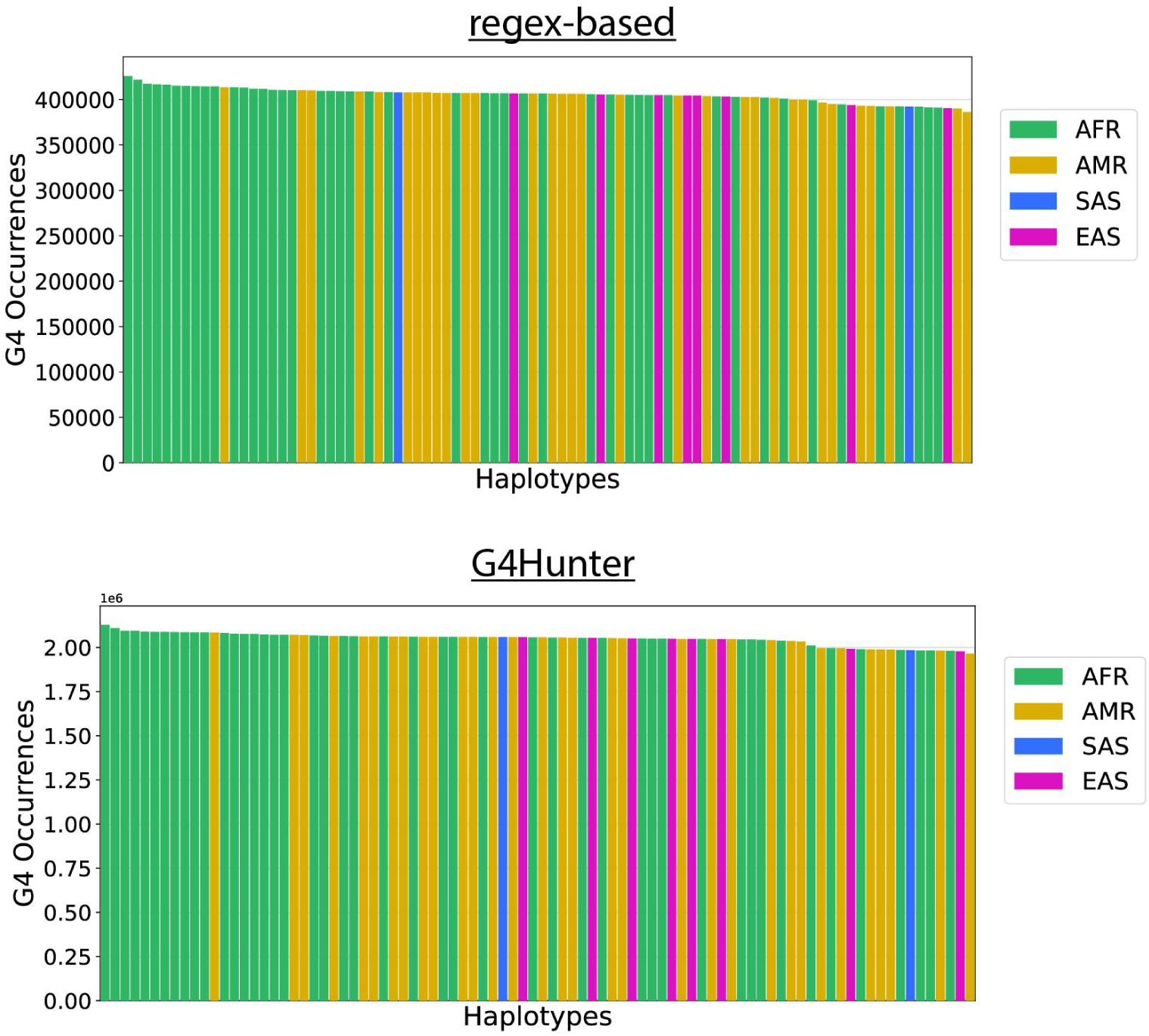
Total G4 occurrences per haplotype. Top panel represents G4s from the regex-based algorithm, and the bottom panel from the G4Hunter algorithm. Colors represent the ancestry of each haplotype.

**Supplementary Figure 2:**
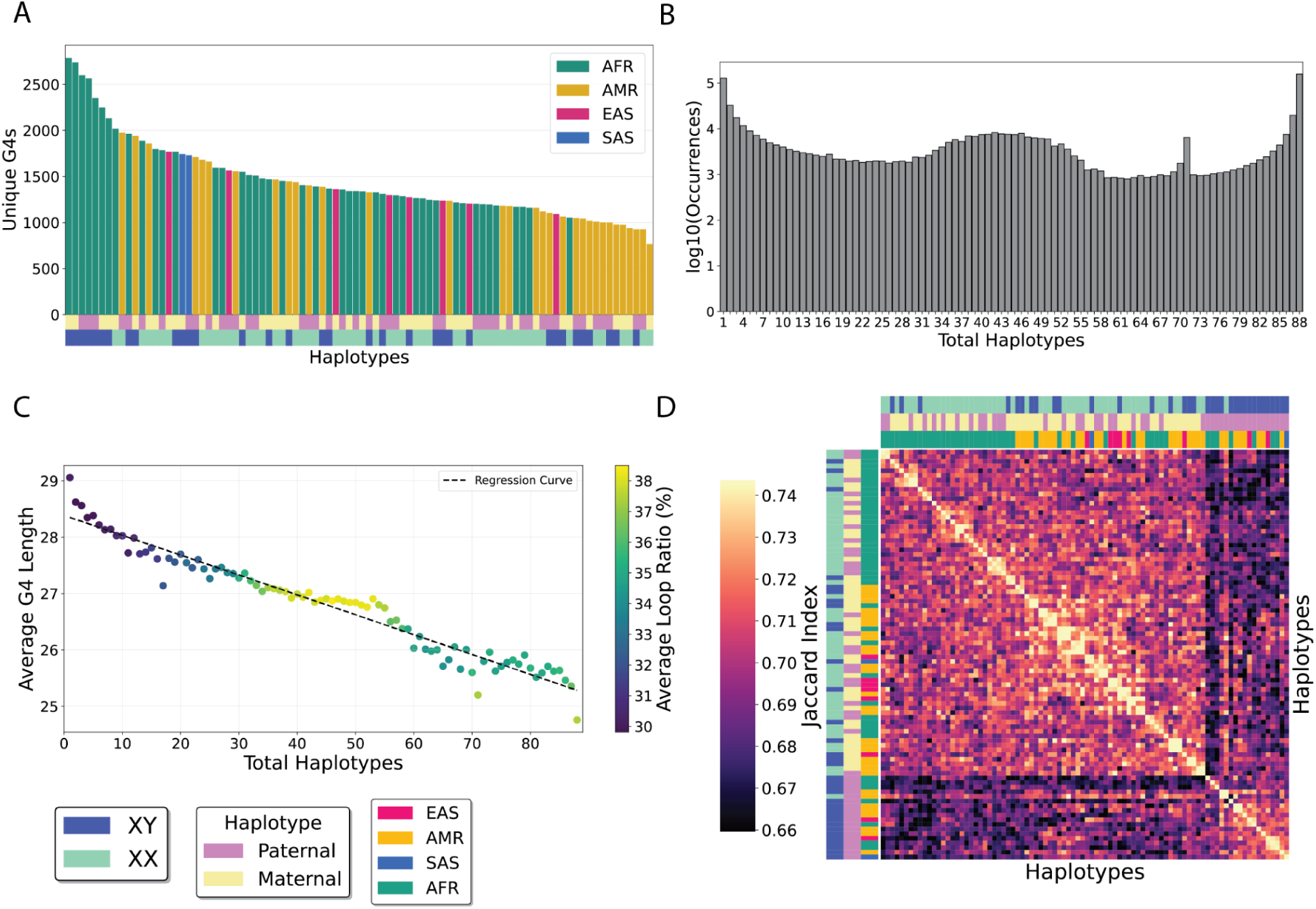
Characterization of G4s across the T2T reference human genome using the regex-based algorithm. **A.** Number of unique G4s per human haplotype. **B.** Number of G4s motifs uniquely shared across different numbers of haplotypes. **C.** Number of G4 motifs uniquely shared across different numbers of haplotypes plotted against average G4 length. **D.** Hierarchical clustering of conserved G4s found in the reference genome CHM13v2 across 88 haplotypes.

**Supplementary Figure 3:**
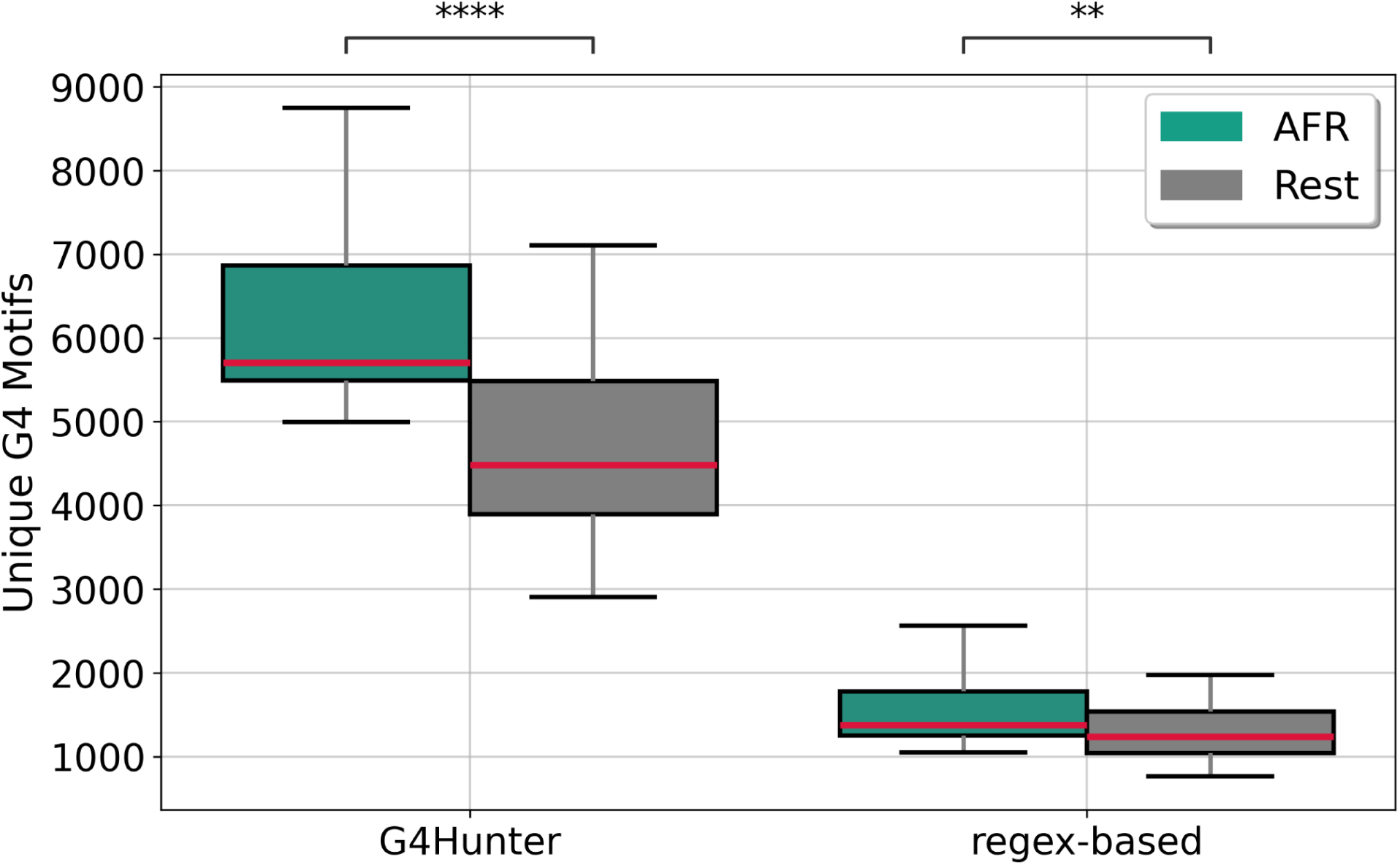
Comparison of the number of unique G4 motif sequences identified in AFR haplotypes versus all other haplotypes, using both G4Hunter and regex-based methods. Independent two-tailed t-tests were performed, with p-values adjusted for multiple comparisons using the Bonferroni correction.

**Supplementary Figure 4:**
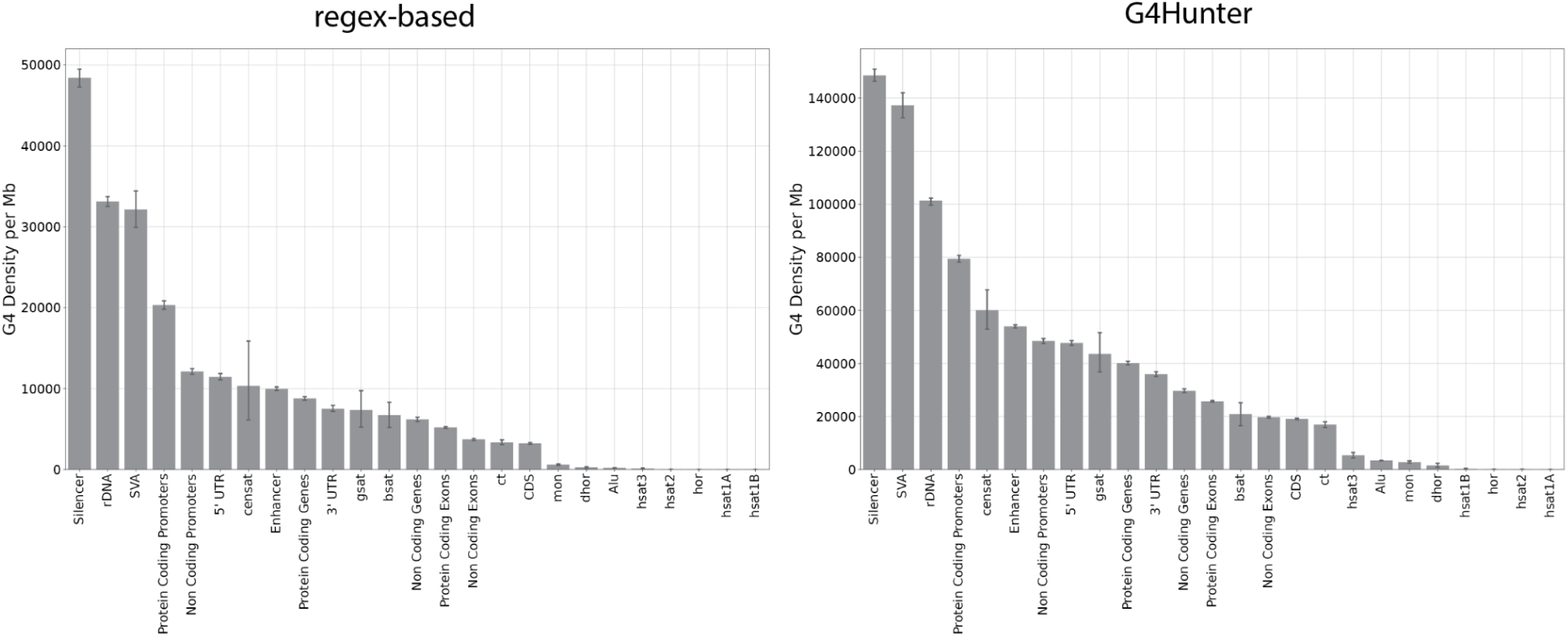
G4 density per Mb across various genomic subcompartments for both G4Hunter and regex-based G4s. Telomeres were not included in the diagrams.

**Supplementary Figure 5:**
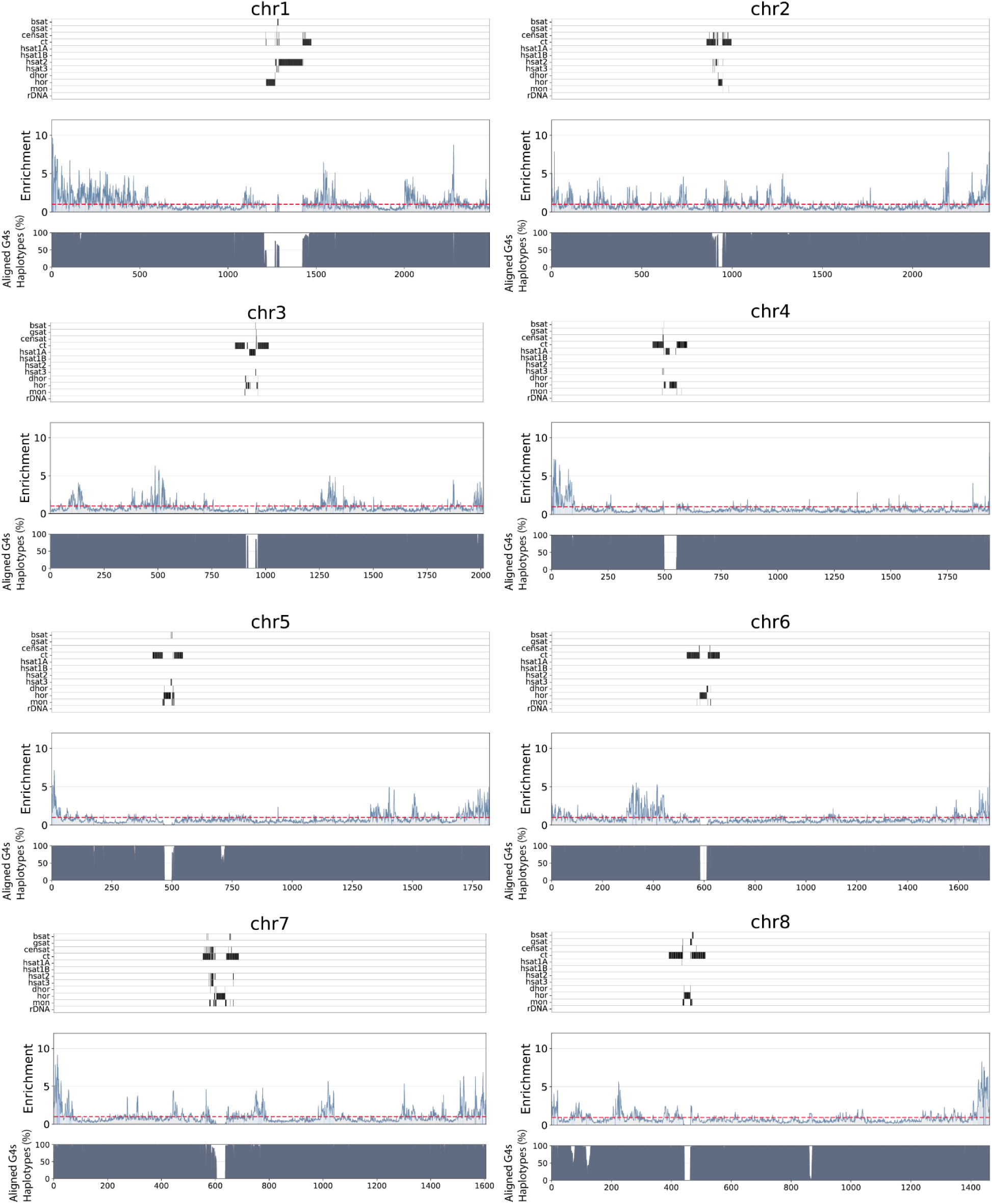
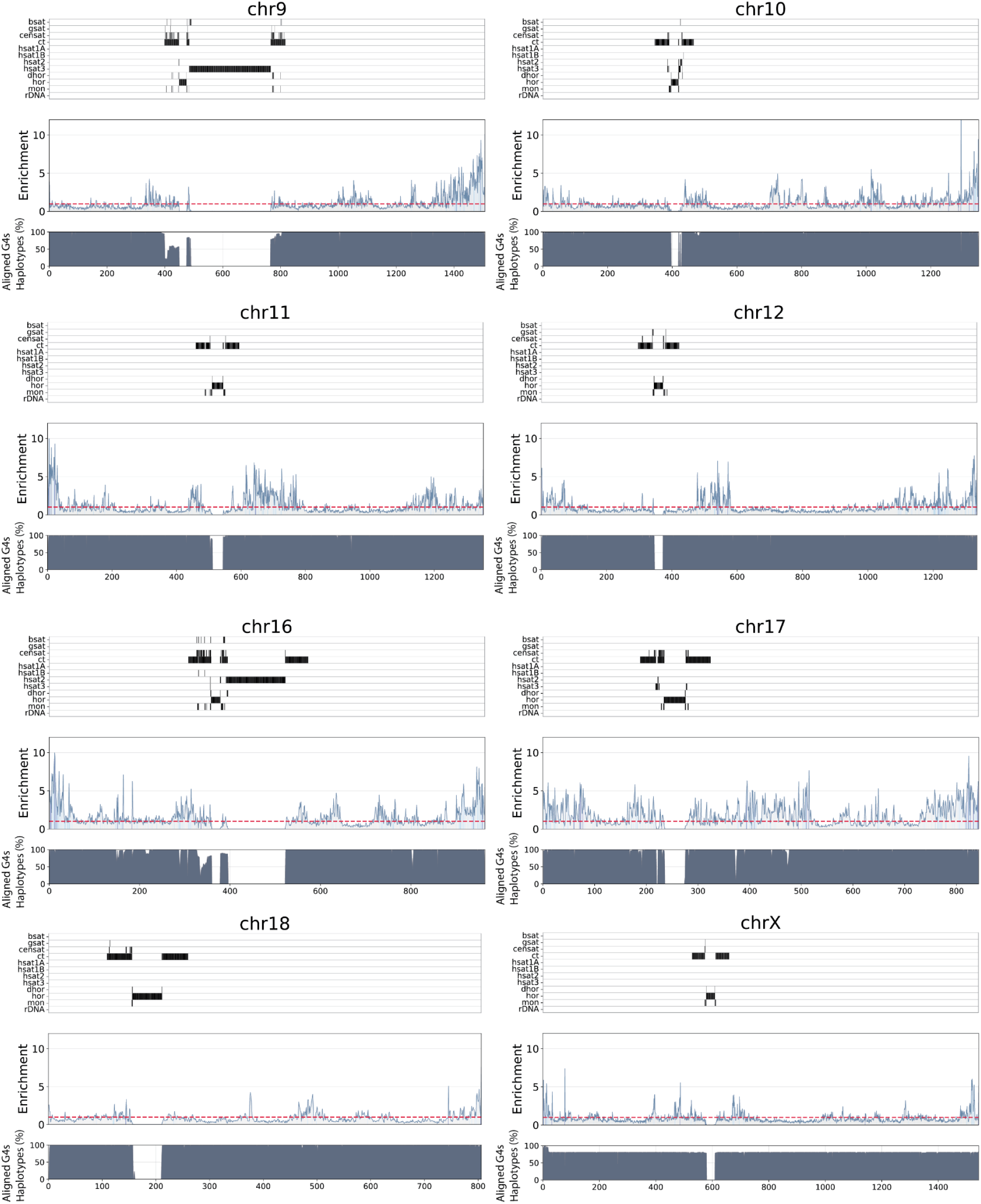
Enrichment of G4s across human chromosomes, each one equipartitioned into 2,000 mutually exclusive regions. The area under the curve denotes the significance of the enrichment, with green representing a significant enrichment, and red a non-significant enrichment, when adjusting for GC-content. The top panel aligns the position of centromeric and pericentromeric regions. The bottom panel illustrates, for each bin, the average percentage of haplotypes for which aligned G4s were conserved in a given bin. In a given bin, the white stripes represent either the absence of G4s or a low percentage of haplotypes. Repeats include inactive αSat HOR (hor), divergent αSat HOR (dhor), monomeric αSat (mon), classical human satellite 1A (hsat1A), classical human satellite 1B (hsat1B), classical human satellite 2 (hsat2), classical human satellite 3 (hsat3), β-satellite (bsat), γ-satellite (gsat), other centromeric satellites (CenSat), and centromeric transition regions (ct).

**Supplementary Figure 6:**
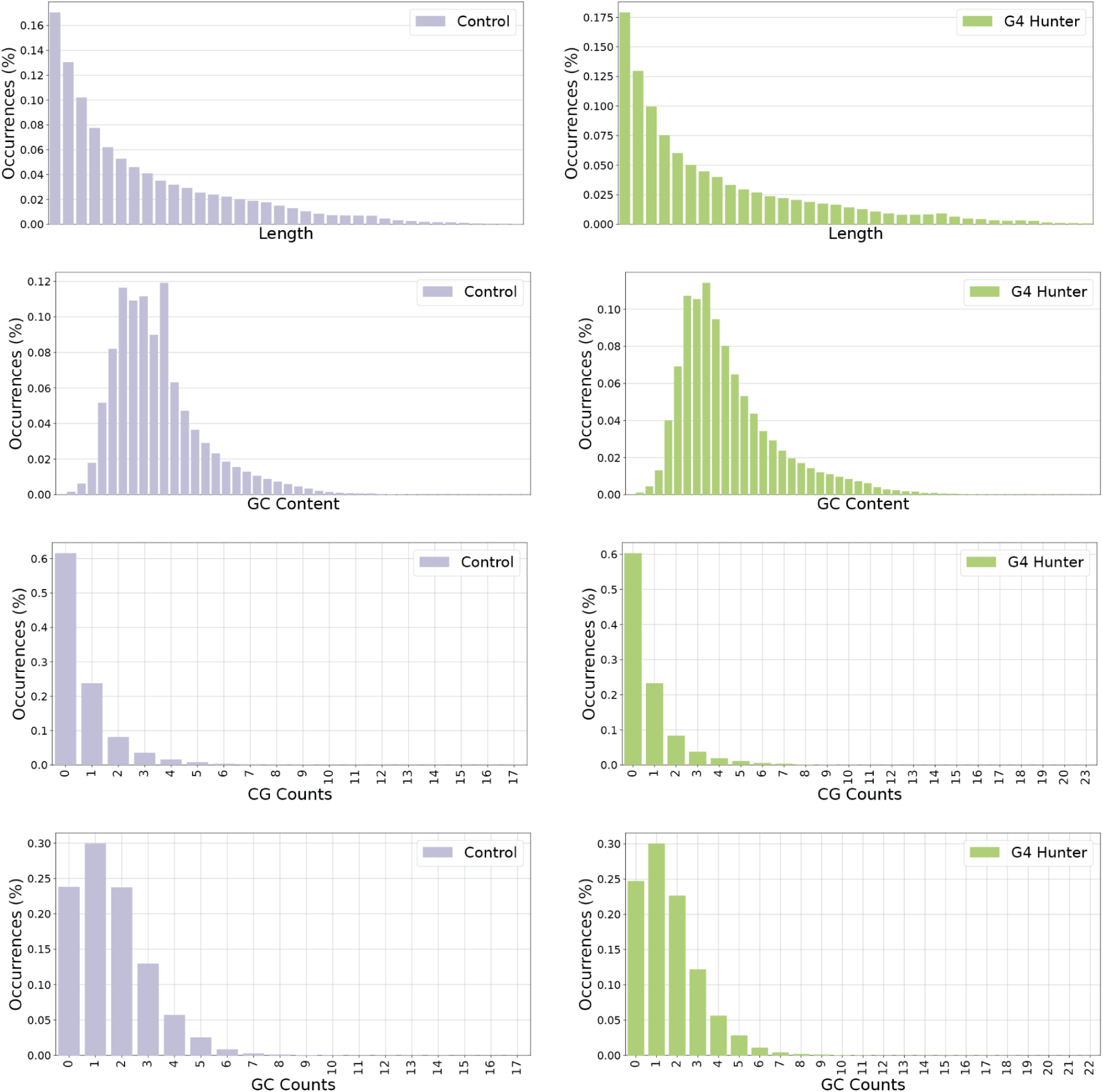
Nucleotide composition comparison between G4s from G4Hunter and G4 controls.

**Supplementary Figure 7:**
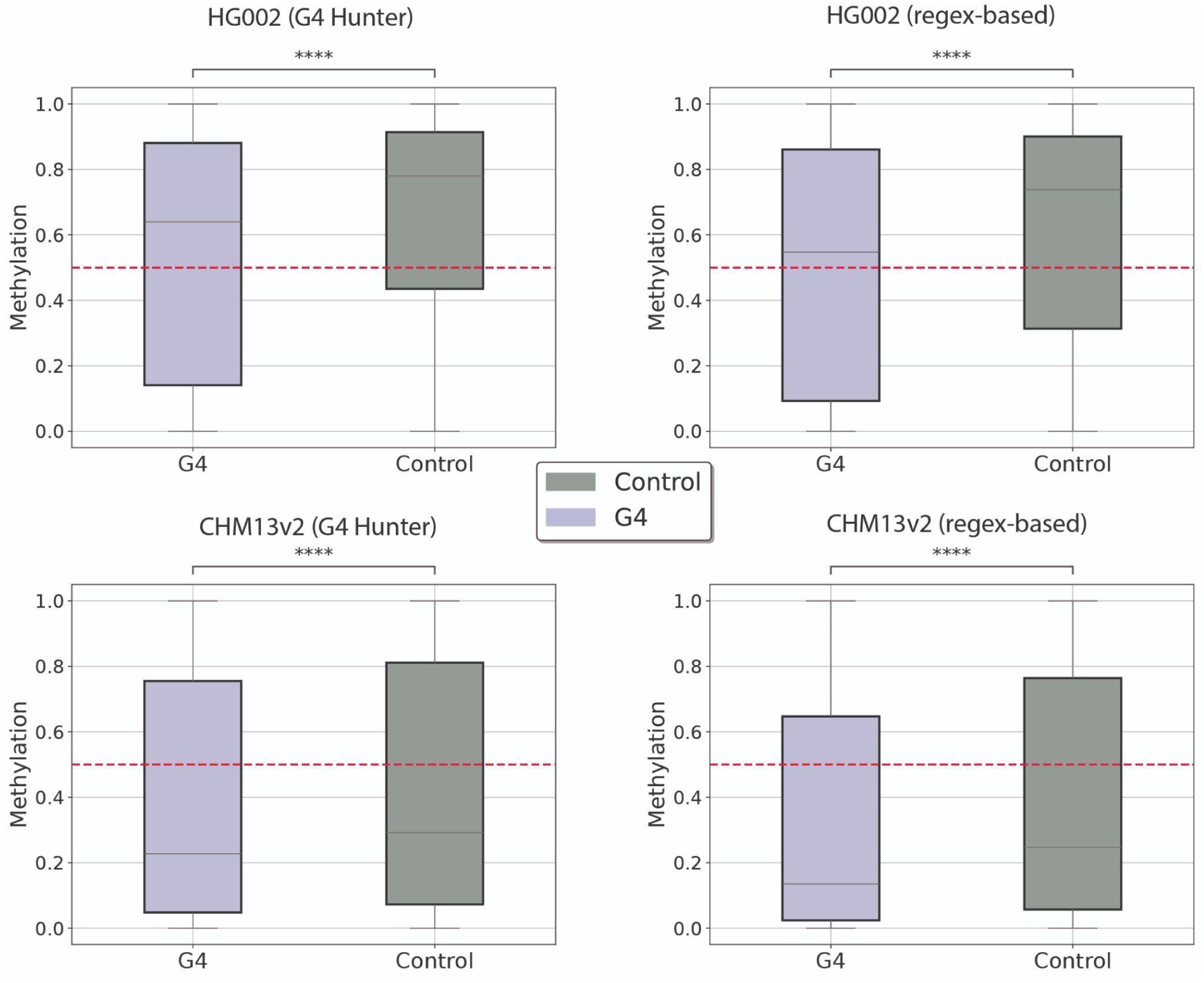
Comparisons of control group and G4 for both regex-based and G4Hunter extraction algorithms for genome-wide methylation profiles across lymphoblastoid and complete hydatidiform mold dell lines. Results shown using Mann-Whitney U tests.

**Supplementary Figure 8:**
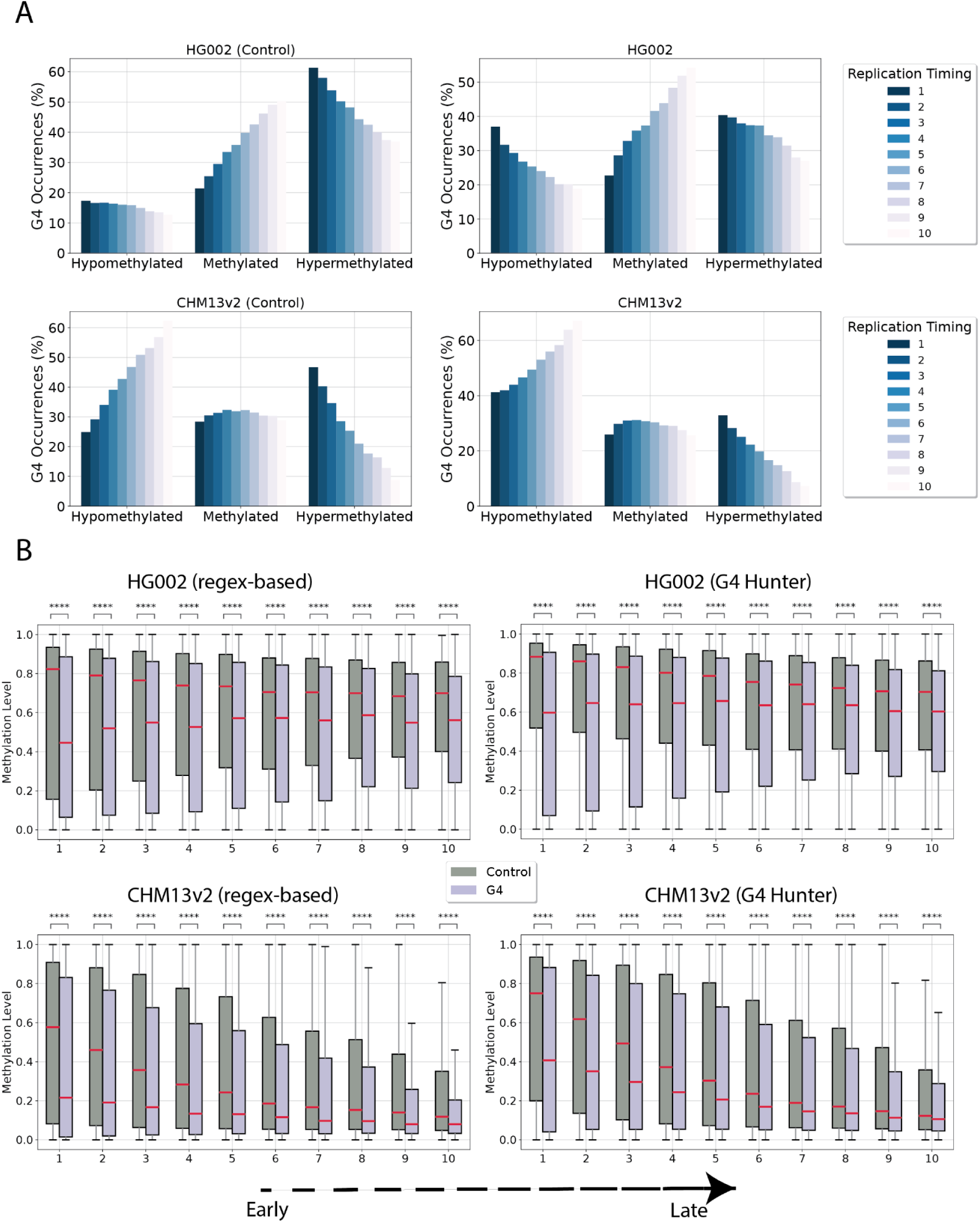
Replication timing and G4 methylation across two different cell lines. **A.** Percentage of hypomethylated, methylated, hypermethylated controls and G4s across lymphoblastoid (HG002) and complete hydatidiform mold (CHM) cells for decreasing replication timing. **B.** Comparisons of control group and G4 for both regex-based and G4Hunter extraction algorithms for decreasing replication timing for the lymphoblastoid and the CHM methylation patterns. Bonferonni adjusted comparisons.

**Supplementary Figure 9:**
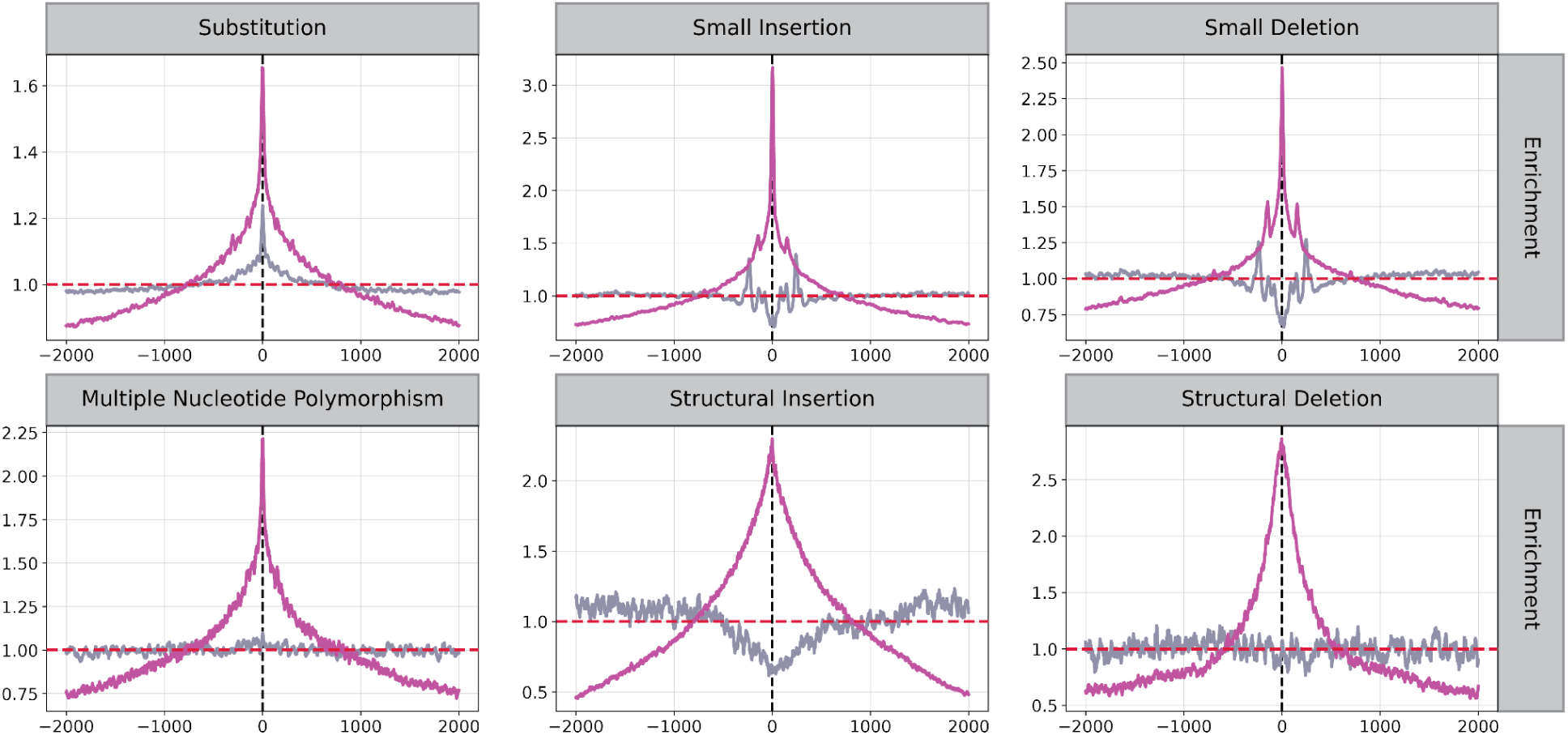
Relative positioning of regex-based G4s (in magenta) and control group (gray) across various mutation loci.

**Supplementary Figure 10:**
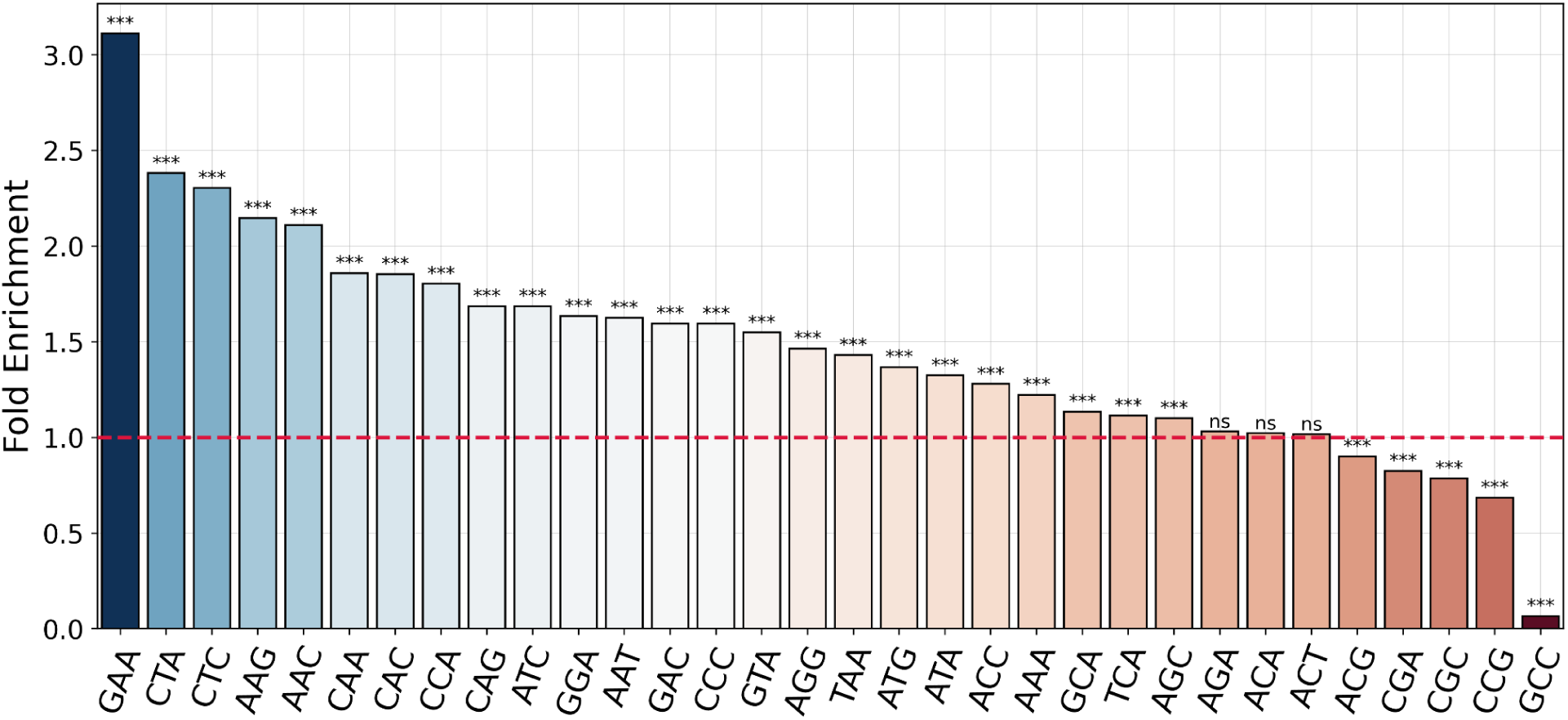
Trinucleotide substitution model showing which trinucleotides are more frequently mutated at G4 motifs compared to matched control regions. Τrinucleotides refer to a mutated base and its immediate 5’ and 3’ neighboring bases. This captures the influence of local sequence context on substitutions at G4s. Significance of the enrichment has been assessed using a two-tailed binomial test.

**Supplementary Figure 11:**
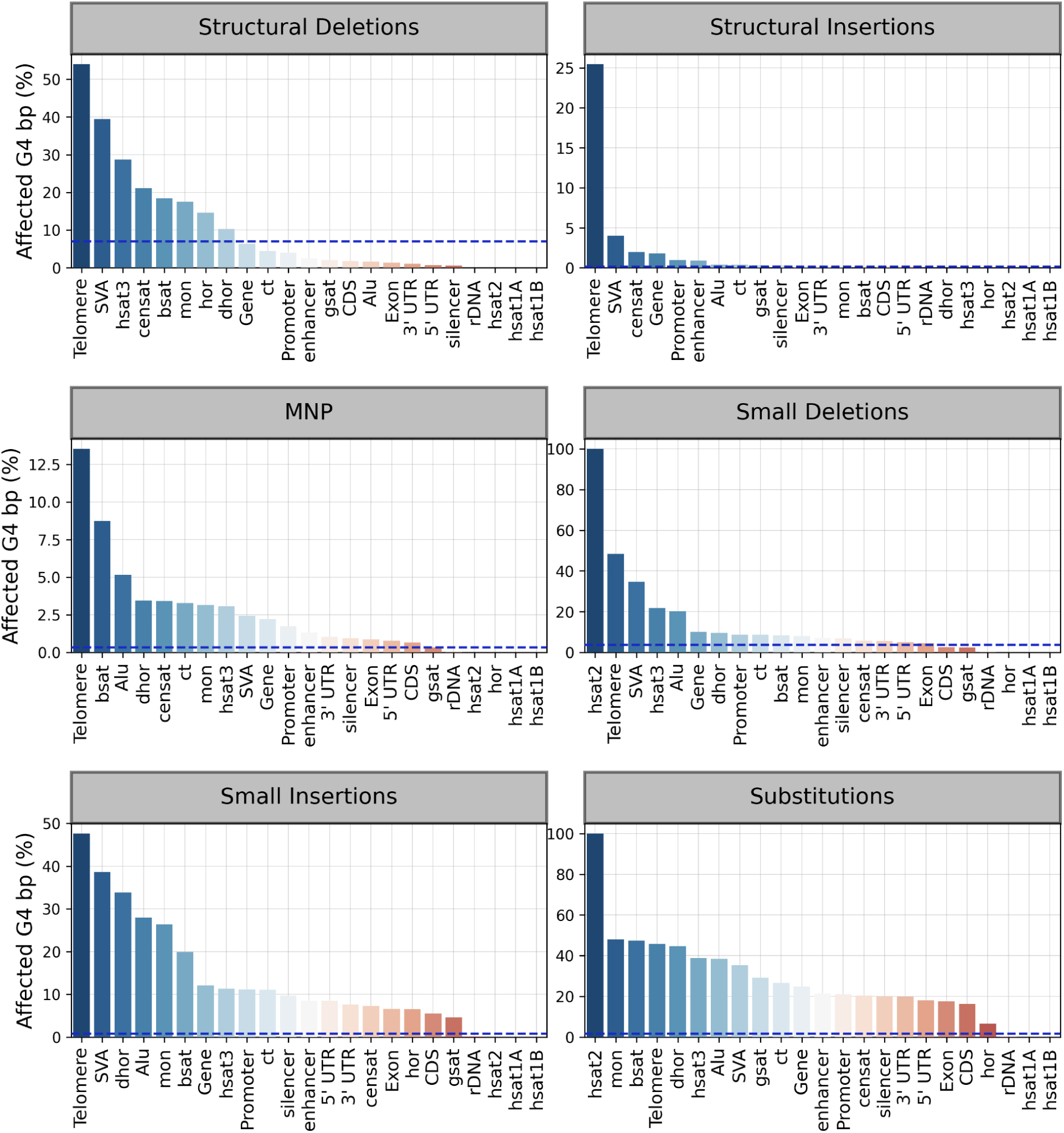
Mutation rate of regex-based G4 base pairs across various genomic subcompartments of interest. In the Human Pangenome Reference Consortium, mutations falling in rDNA loci in many cases were masked.

**Supplementary Figure 12:**
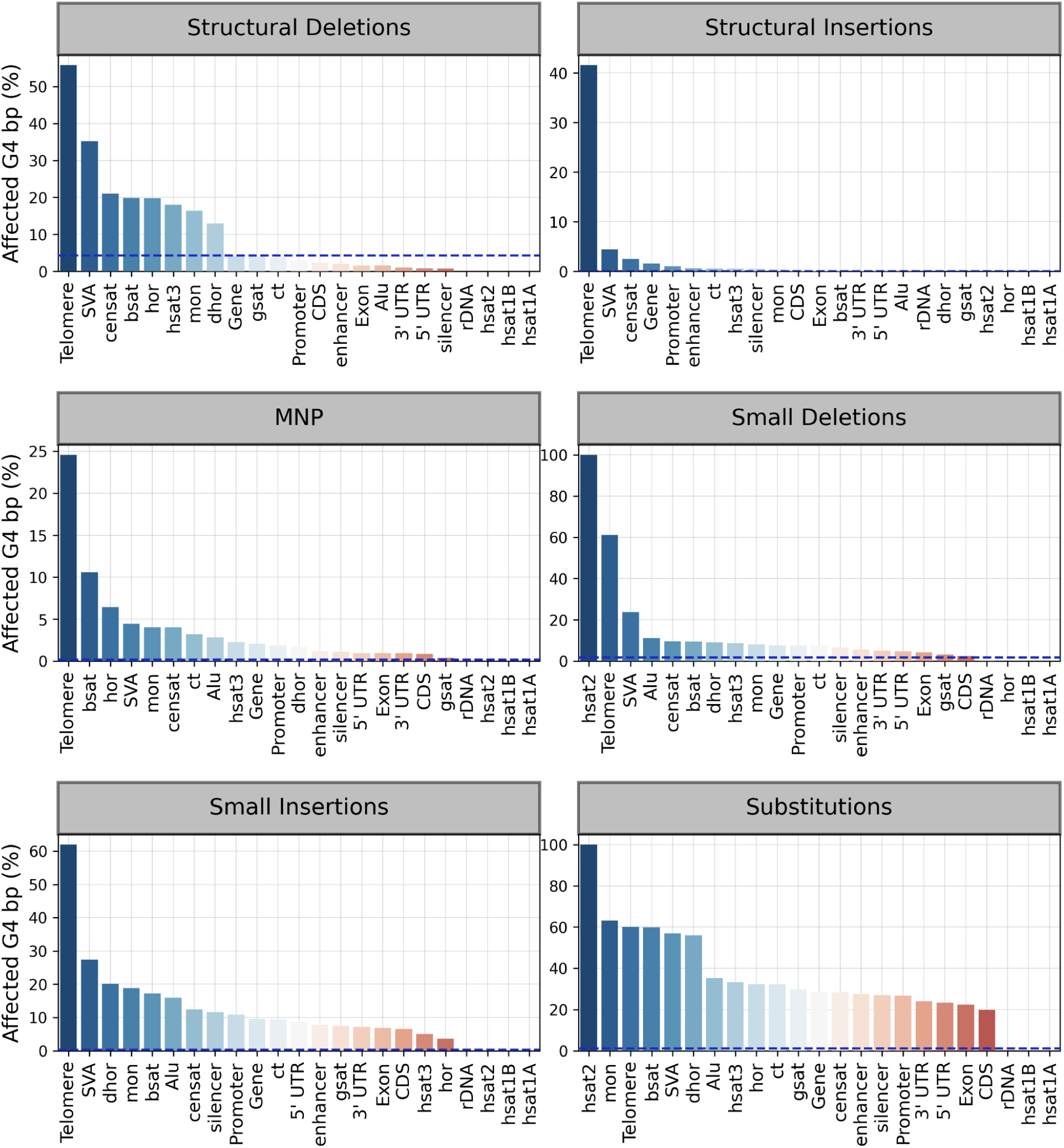
Mutation rates of G4Hunter extracted G4 base pairs across various genomic subcompartments of interest. In the Human Pangenome Reference Consortium, mutations falling in rDNA loci in many cases were masked.

**Supplementary Figure 13:**
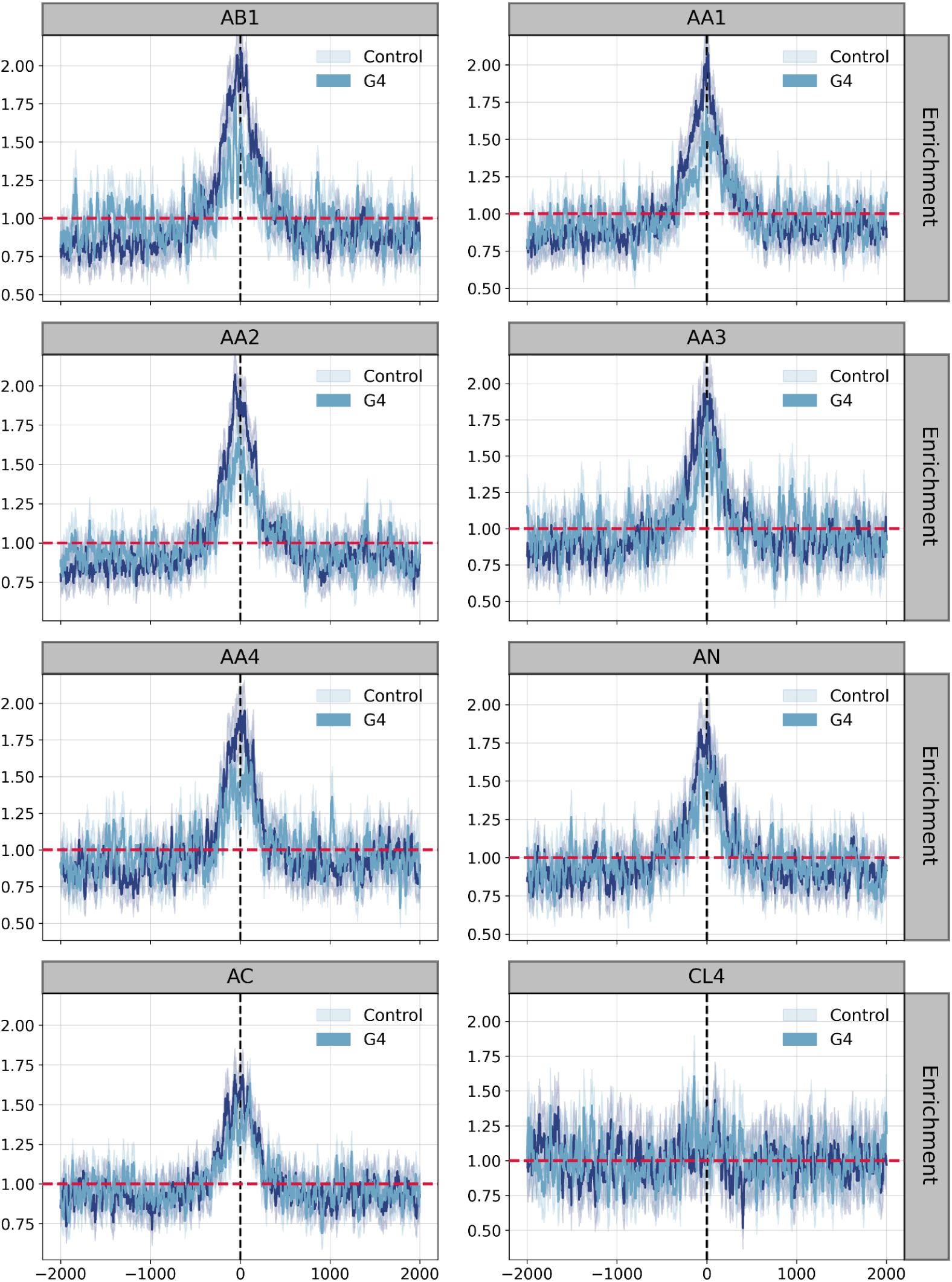
PRDM9 regex-based algorithm 2kB density plot relative to PRDM9 loci for various alleles.

**Supplementary Table 1:**
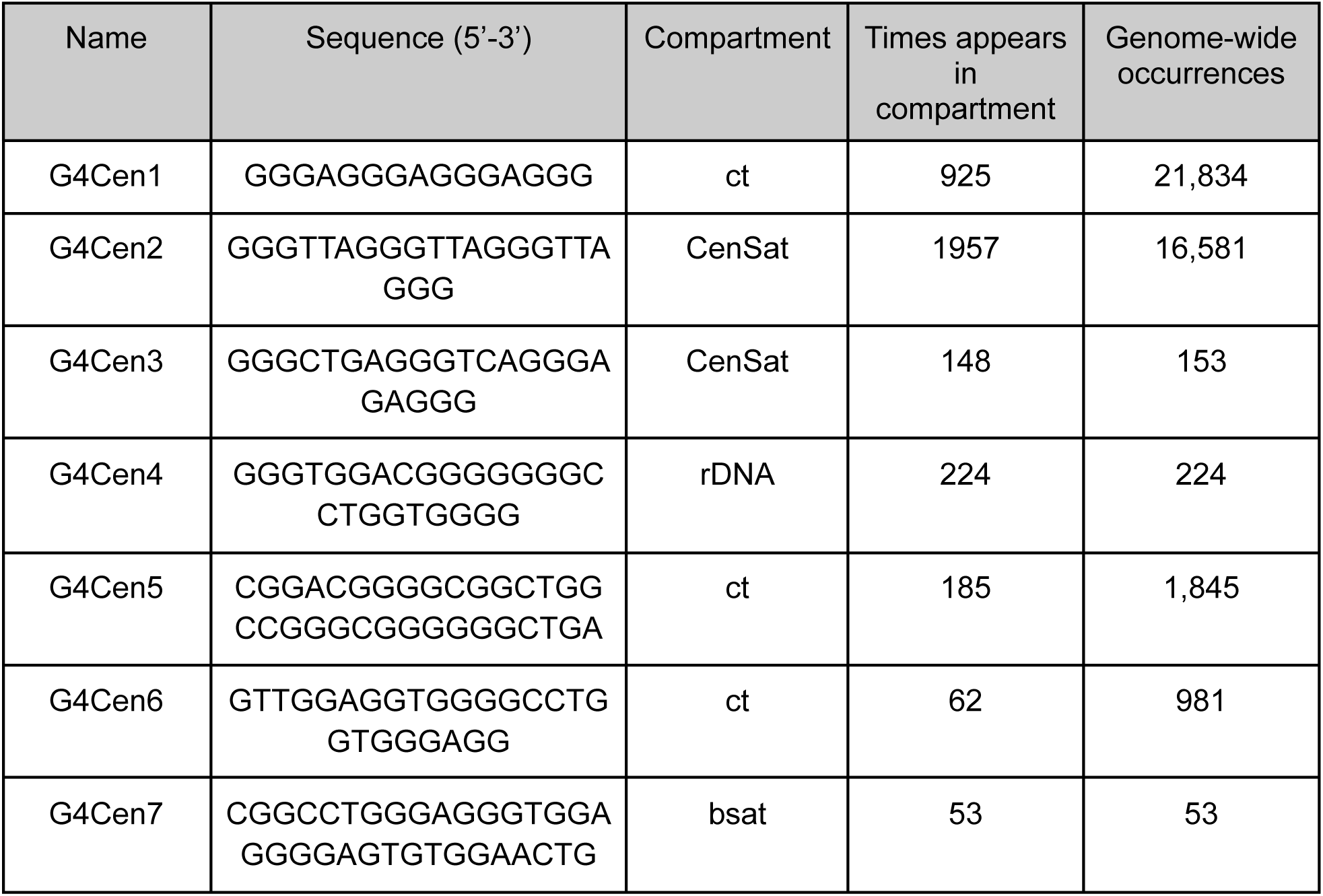
Most frequent canonical G4s within centromeric, pericentromeric and satellite regions. Selected sequences found in highly repetitive parts of the human genome based on G4 consensus and G4Hunter. Occurrences represent the genome-wide counts.

## Notes

https://github.com/Georgakopoulos-Soares-lab/g4_t2t_identification

### Competing Interest Statement

The authors have declared no competing interest.

### Summary of Updates

Author name typo fixed; Updated GitHub link.

